# Myeloid cell protein tyrosine phosphatase 1B (PTP1B) drives retinal neurodegeneration in diabetic mice

**DOI:** 10.1101/2025.10.15.682409

**Authors:** Ayaz Ali, Morgan Boyne, Abrar Othman, Sarah Kamli-Salino, Lucia Kuffova, John V. Forrester, Mirela Delibegovic

## Abstract

Diabetic retinopathy (DR) is the leading cause of vision loss in the working age population with public health economic implications worldwide. Systemic inflammation and leukocyte activation are early events in diabetes, while microglial activation, neuroinflammation, and retinal neurodegeneration are early events in DR. Protein tyrosine phosphatase 1B (PTP1B) plays a complex role in monocyte / macrophage activation which may impact DR. We therefore investigated the role of myeloid cell-specific PTP1B using LysM^cre^-PTP1B fl/fl (LysM-PTP1B) mice, as well as a PTP1B inhibitor, MSI-1436, in the early stages of DR. Mice were rendered diabetic for six weeks using anomer-equilibrated streptozotocin (STZ). Retinal changes were evaluated by histology and immunohistochemistry, and systemic leukocyte activation by flow cytometry. Mitochondrial function in high glucose-challenged, cultured bone marrow-derived macrophages (BMDMs) from LysM-PTP1B and MSI-I436-treated mice was determined *in vitro*. Both myeloid cell-specific depletion and pharmacological inhibition of PTP1B prevent STZ-induced retinal neurodegeneration, development of acellular retinal capillaries, as well as microglial and systemic leukocyte activation without effect on development of diabetes. *In vitro*, inhibition of PTP1B prevented high glucose-induced mitochondrial dysfunction in BMDMs. We conclude that inhibition of PTP1B prevents DR by decreasing myeloid cell-driven inflammation and PTP1B represents a therapeutic target for prevention DR.

**Highlights:**

- Myeloid cell PTP1B is required to induce retinal neurodegeneration in diabetes
- Both local (microglia) and systemic (bone-marrow derived) myeloid cells are implicated
- Inhibition of myeloid cell PTP1B prevents development of acellular retinal capillaries in diabetic mice
- PTP1B mediates superoxide production, decreases mitochondrial membrane potential and increases macrophages cell death in chronic conditions associated with abnormally high glucose.

**Graphical abstract:** 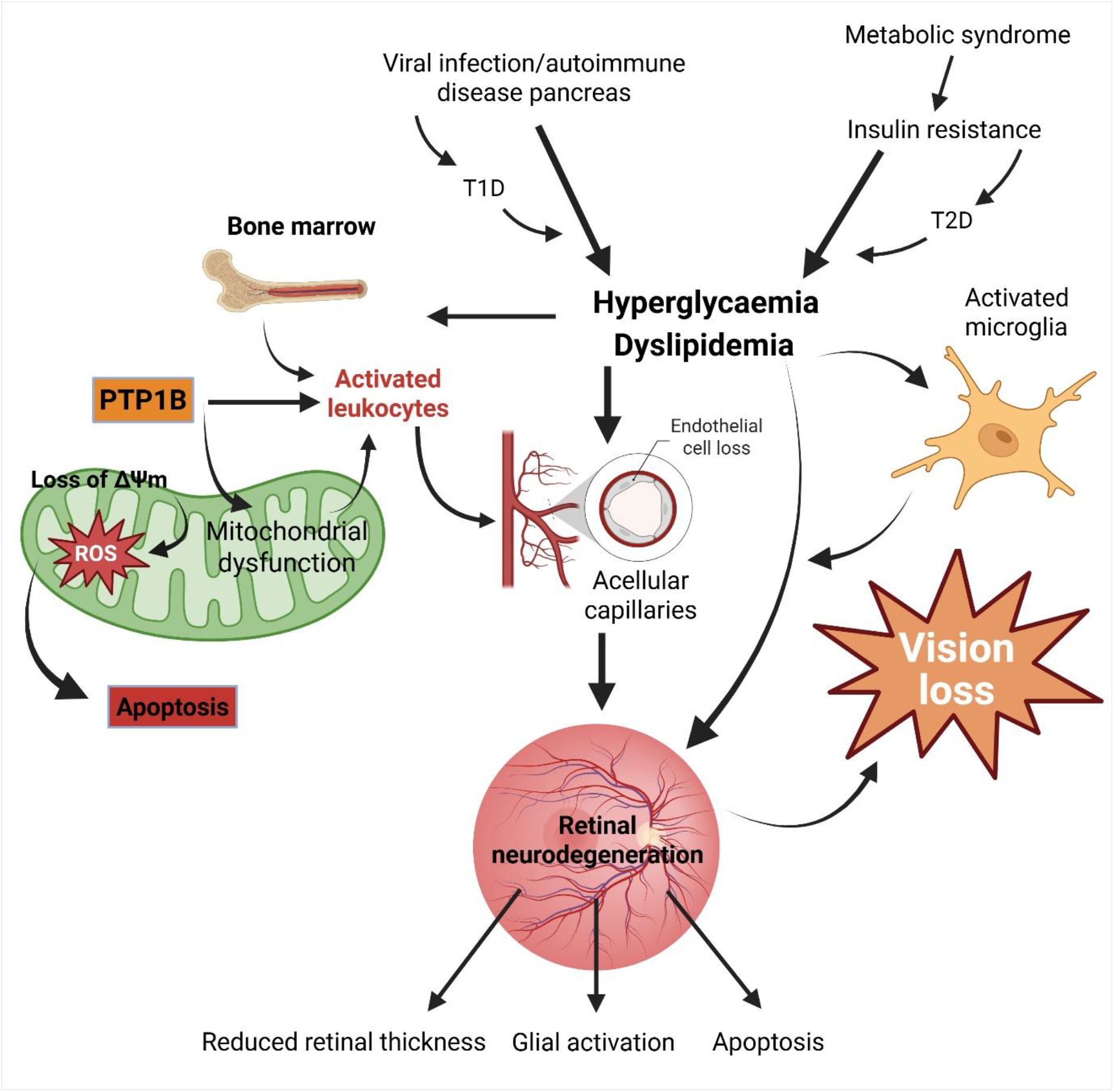

## Introduction

Diabetic retinopathy (DR) is the leading cause of sight-threatening disease in the working age population with serious health economic implications (1). DR is characterised by dysfunction of the retinal neurovascular unit (NVU) and other microvascular abnormalities in response to prolonged hyperglycaemia in type 1 and type 2 diabetes (2,3). Non-proliferative DR presents with increased retinal vascular permeability, leukostasis, capillary occlusion, retinal ischemia and neuronal degeneration during the early stages of diabetes and can later progress to proliferative DR and consecutive loss of vision (4). Thus, there is urgency to better understand how the early events occur in DR.

Retinal neurodegeneration is an early sign of DR (5). Poorly controlled blood glucose is associated with retinal ganglion cell loss and retinal thinning, with glial cell activation (6) (primarily microglia), and release of pro-inflammatory cytokines and chemokines (7). However, the underlying mechanisms for early neurodegeneration and neuroinflammation in diabetes are not fully understood.

We and others have shown that leukostasis is one of the earliest signs of DR in the streptozotocin (STZ) mouse model of diabetes and DR, and is caused by CD11b+CCR5+ myeloid cells trapped in retinal capillaries (8) leading to capillary occlusion, loss of capillaries, vascular leakage and retinal ischemia (9). Moreover, inhibition of leukocyte adhesion prevents the development of DR (3). Low-level myeloid and T cell activation (10–12) establishes a background of systemic inflammation (parainflammation) (10,13) due to disturbed glucose and lipid metabolism. Hyperglycaemia itself is directly proinflammatory, as shown by an increase in resident tissue macrophages in several tissues, which promotes increased neutrophil recruitment after challenge (14). Interestingly, a recent study demonstrated that increased CC chemokine receptor 2 (CCR2) derived inflammatory monocytes contribute to the retinal capillary degeneration in the early stages of DR in mice (15). Further work has shown that CCR2 mediates the mobilisation and migration of bone marrow-derived classical monocytes to injury sites including the retina in diabetes (16–18) while pharmacological inhibition of CCR2/5 prevents STZ-induced increased vascular permeability in diabetic mice (19). These studies suggest that the majority of leukocytes involved in the vasculopathy DR are systemically (bone marrow) derived. However, whether the neurodegeneration is due to early changes in leukocyte activation (i.e. inflammation) or is it a secondary effect of the metabolic changes (e.g. high blood glucose) on retinal cells remains an unsolved question of practical clinical importance.

Protein tyrosine phosphatase 1B (PTP1B), a non-receptor tyrosine phosphatase with pleiotropic roles in both the immune response (20) and glucose metabolism (21), is a negative regulator of insulin and leptin receptors (22), and a tumour suppressor (23). Global PTP1B depletion in mice ameliorates diet-induced obesity and prevents insulin resistance (24,25). Myeloid cell-specific PTP1B deletion in mice challenged with high dose lipopolysaccharide and D-galactosamine (26) generates increased susceptibility to the effects of irradiation (27) suggesting a crucial role for macrophage-specific PTP1B in regulating immune responses. Our earlier studies demonstrated that depletion of PTP1B in macrophage/monocytes prevents inflammation-driven pathology in obesity-associated insulin resistance in mice (28,29). However, the role of myeloid cell-specific PTP1B in the complications of STZ-induced diabetes, particularly in DR, is unknown.

In the current study, we investigated the role of myeloid cell-specific PTP1B in the early stages of retinal neurodegeneration, utilising myeloid cell PTP1B-depleted (LysM-PTP1B) STZ-induced diabetic mice. In parallel, we investigated the therapeutic effects of the small molecule PTP1B inhibitor, MSI-1436, on retinal neurodegeneration in STZ-induced diabetic C57BL/6J mice. Our results demonstrate that both, myeloid-specific deletion and pharmacological inhibition of PTP1B, prevent STZ-induced retinal neurodegeneration, highlighting PTP1B as possible therapeutic target for the treatment and prevention of diabetic retinopathy.

## Research Design and Methods

### Mice and mouse husbandry

The animal procedures used in this study were approved by the UK Home Office Animals (Scientific Procedures) 1986 Act (PPL PP3110164) and in accordance with the Association for Research in Vision and Ophthalmology (ARVO) statement for the use of animals in vision and ophthalmology research. The generation of myeloid cell PTP1B-depleted mice on the C57BL/6J background was described previously (30) (additional details in the Supplementary Text).

### Induction of STZ-induced diabetes model and study design

Diabetes was induced in sixteen- and twenty-four-week-old male and female PTP1B fl/fl and LysM-PTP1B mice using 40 and 50 mg/kg respectively of anomer equilibrated streptozotocin (STZ) (Sigma-Aldrich, USA) as described previously (31) (Supplementary Fig. 1). For the PTP1B inhibitor treated model, ten-week-old non-diabetic and diabetic mice were injected intraperitoneally with the PTP1B-specific allosteric inhibitor, MSI-1436 (1 mg/kg) dissolved in saline, whereas the vehicle group was administered with saline once per week for eight consecutive weeks (additional details in the supplementary Text).

### Body composition measurement

Body composition (total fat and lean mass) was assessed using a magnetic resonance imaging 3-in-1 scanner (Echo, Medical Systems, Houston, USA) (additional details in the Supplementary Text).

### Flow cytometry of spleen and bone marrow (BM)

Single cell suspensions from the mouse BM and spleen tissues were prepared as described previously (8). Tissues were mechanically dissociated with a 20 ml syringe plunger and passed through a 70 µm cell strainer, washed and subsequently resuspended in RBC lysis buffer (Sigma-Aldrich, USA) for 5 mins. Thereafter, cells were washed and incubated with the fixable viability dye, eFluor 450, (Biolegend, UK) at 1:1000 dilution. Fc receptors were blocked using CD16/32 (clone 2.4G2) antibody (1 µl/1 × 10^6^ cells) for 30 min at 4 ℃. Cells washed in PBS were labelled with the fluorochrome conjugated surface monoclonal antibodies in FACS buffer [(0.5% bovine serum albumin (BSA), and 2 mM EDTA in PBS (Gibco, Fisher Scientific, UK)] for 30 min at 4 ℃. For individual markers of interest, gating was based on fluorescence minus one (FMO) or isotype controls. Compensation was performed using UltraComp Beads (Invitrogen, UK) and fluorochrome conjugated monoclonal antibodies. A total of 2 × 10^5^ events were acquired on a BD LSR Fortessa flow cytometer (BD Bioscience, UK) and analysed using FlowJo (additional details in the Supplementary Text).

### Retinal thickness measurement by hematoxylin and eosin (H&E) staining

Enucleated eyes fixed in 2.5% glutaraldehyde (Fisher Chemicals, UK) for 48 h were resin embedded, sectioned in 4 µm thick slices, and stained with H&E to evaluate retinal thickness. A total of 7 sections per retina were prepared, each cut at 20 µm intervals distant from the optic nerve and scanned using a Zeiss Axioscan Z1 slide scanner (Carl Zeiss AG, Oberkochen, DE). Three different locations were employed to assess retinal thickness, including posterior pole at 500 µm distance from the optic nerve and at 500 µm distance from the far retinal periphery, while retinal thickness at the mid periphery was measured at a 45º angle from the midpoint of the retina (Supplementary Fig. 2). Retinal thickness was quantified by calculating average of the 7 sections per retina at the three designated regions (posterior pole, mid periphery, and far periphery) from inner limiting membrane to the surface of retinal pigment epithelium at both superior and inferior regions using QuPath software version 0.3.2.

### Immunohistochemistry staining of glial fibrillary acidic protein (GFAP)

Immunohistochemistry staining was performed as stated previously (32). Additional details are provided in the Supplementary Text.

### Terminal deoxynucleotidyl transferase dUTP nick end labelling (TUNEL) assay

TUNEL assay was performed to evaluate apoptotic cell death following the manufacturer’s instruction. Additional details are provided in the Supplementary Text.

### Immunofluorescence staining of microglia

OCT embedded retinal tissues stored at -20°C were defrosted and fixed in 4% methanol-free paraformaldehyde for 30 min. Next, retinal tissues were incubated with the blocking solution containing 1% BSA in TBS-0.2% Triton X-100 (Sigma-Aldrich, US) for 2 h at 4°C. Further, tissues were probed with the ionised calcium binding adapter molecule 1 (Iba-1) primary antibody (Invitrogen, UK) at 1:200 dilution overnight at 4°C. Retinal tissues were washed and subsequently incubated with Alexa Fluor 555 secondary antibody (Invitrogen, UK) using 1:200 dilution for 2 h at 4°C and counterstained with DAPI. Lastly, tissues were mounted, and images were captured using fluorescence microscopy (Carl Zeiss AG, Oberkochen, DE). Retinal microglia morphology was assessed following a previously described method (33). Retinal images were converted into binary and skeletonised images to analyse the morphology ranging from ramified to amoeboid morphology. Parameters such as number of Iba-1+ cells per field, number of microglial process endpoints and the process length of individual cells were measured using Analyze Skeleton plugin in Fiji ImageJ image analysis software.

### Detection of endothelial cell loss and acellular capillaries in retinal flat mounts

Retinas from eyes fixed in 4% paraformaldehyde for 2 h were flatmounted and fixed in ice cold methanol for 20 min. The flatmounts were then blocked in blocking solution (2% BSA in PBS - 0.3% Triton X-100) for 2h at RT followed by incubation with primary antibodies, isolectin B4 (Vector Laboratories, B-1205, UK) and collagen IV (Bio-Rad, 2150-1470, UK) at 1:100 dilution in blocking solution overnight at 4°C. Next, retinal flatmounts were probed with secondary antibodies, streptavidin Alexa Fluor 488 (S32354, Invitrogen, UK) and donkey anti-rabbit Alexa Fluor 594 (Jackson ImmunoResearch, 711-585-152, UK) at 1:300 dilution for 2h at RT. Finally, retinas were washed and mounted on a glass slide using mounting medium. Images were captured using a confocal microscope (LSM 880, Carl Zeiss AG, DE). Acellular capillaries were identified as collagen IV positive, isolectin B4 negative vessels and their number counted in all three vascular layers (superficial, intermediate, and plexus) of the retina by taking average of the 3-5 images per retina. Vascular density was quantified using AngioTool0.6a as described previously (34).

### Isolation and preparation of bone marrow-derived macrophages (BMDMs)

BMDMs were prepared after flushing the femurs and tibias from age and gender matched non-diabetic and diabetic mice as described previously (28). Briefly, isolated cells were cultured in normal (5 mM) and high glucose (25 mM) medium at 37°C in a humified incubator for 7-8 days supplemented with 10% foetal bovine serum (FBS), L-glutamine and 20% L929 conditioned medium.

### Measurement of mitochondrial membrane potential and superoxide generation

BMDMs cultured in normal and high glucose medium for 7 days were incubated with (a) 5,5’,6,6’-tetrachloro-1,1’,3,3’-tetraethylbenzimidazolylcarbocyanine iodide (JC-1) (Invitrogen, UK) to determine mitochondrial membrane potential and (b) MitoSOX (Invitrogen, UK) reagent to assess mitochondrial superoxide generation following the manufacturer’s guidelines.

### Detection of apoptosis by flow cytometry

BMDMs seeded in normal and high glucose conditions were harvested and incubated with FITC conjugated annexin V and propidium iodide (PI) using binding buffer (BD, Biosciences) for 30 min following the manufacturer’s instructions.

### Statistical analysis

Data are shown as means ± standard deviation (SD). Each n value represents a single mouse in both *in vivo* and *ex vivo* studies. Statistical analyses were conducted employing one way or two-way analysis of variance (ANOVA) followed by Tukey’s or Bonferroni multiple comparison tests and a p value of <0.05 considered statistically significant using GraphPad Prism8 software.

### Data and Resource Availability

“The datasets generated and/or analyzed during the current study are available from the corresponding author upon reasonable request”.

## Results

### Myeloid cell PTP1B is required for STZ-induced retinal neurodegeneration

LysM-PTP1B and PTP1B-floxed control mice (twenty-two- and thirty -week-old) were rendered diabetic using five daily injections of anomer-equilibrated STZ as described in Methods (Supplementary Fig. 1). Data for twenty-two- and thirty -week-old were similar. Results are presented therefore for thirty-week-old mice only unless otherwise stated. Retinal neurodegeneration is now recognised as an early event in diabetes, becoming detectable weeks after the onset of diabetes (35–37). In this study, we measured retinal thickness as described in Methods (Supplementary Fig. 2) and confirmed the early signs of STZ-induced retinal neurodegeneration in male and female mice, namely decreased retinal thickness (Fig. 1A and B), activation of Muller glial cells (increased GFAP expression) (Fig. 2A; Supplementary Fig. 3A) and increased levels of apoptosis (TUNEL staining) (Fig. 2B; Supplementary Fig. 3B).

**Figure 1.**
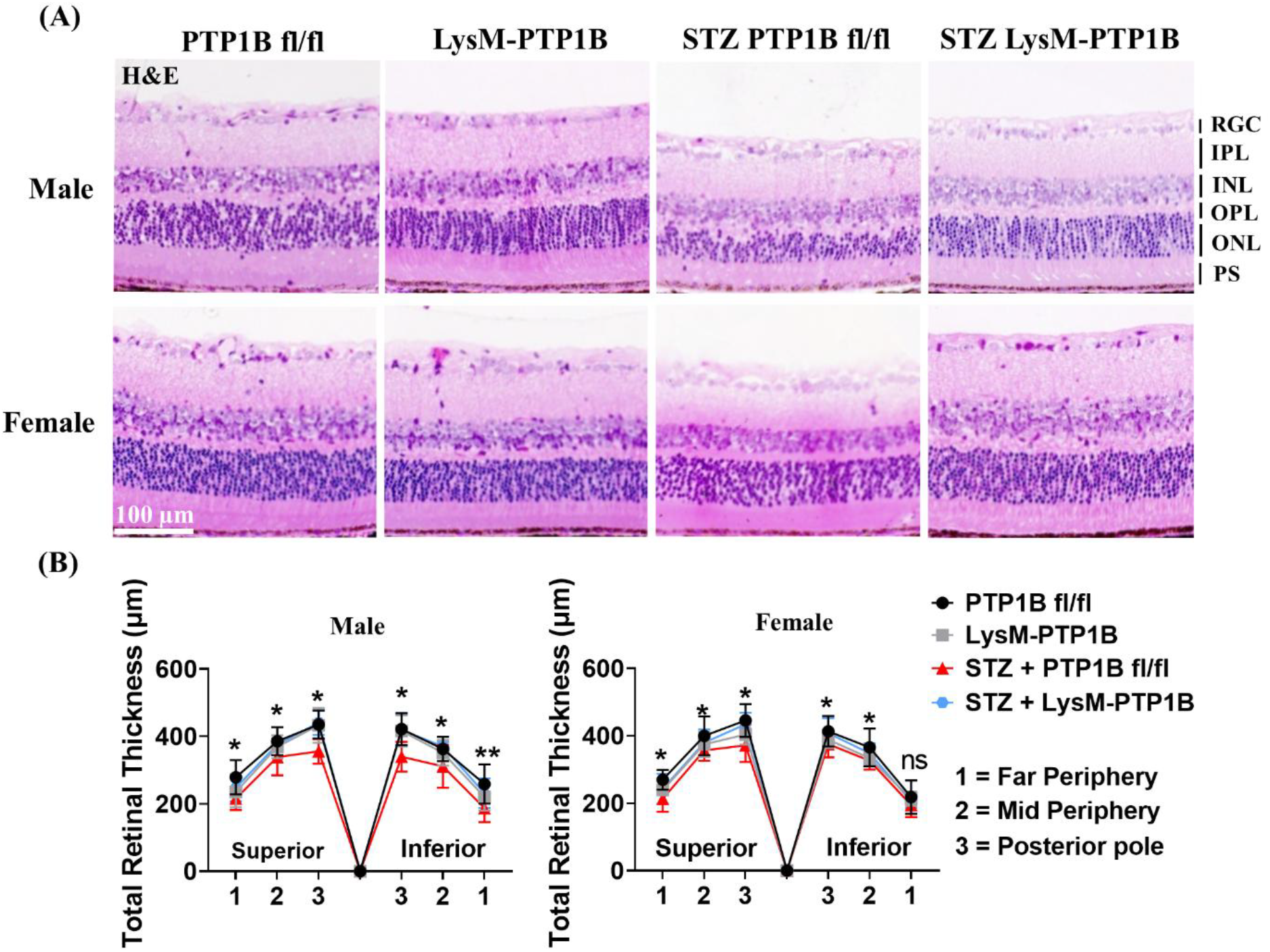
STZ-induced retinal neurodegeneration is prevented in LysM-PTP1B mice. Representative images of resin embedded hematoxylin and eosin (H&E) stained retinal sections of thirty-week-old male and female non-diabetic and diabetic LysM-PTP1B and control littermate PTP1B-floxed mice. Loss of retinal thickness is evident in STZ PTP1B-floxed mice but not STZ LysM-PTP1B mice (A). Scale bar: 100 µm. Measurement of retinal thickness in three different regions of the retina (posterior pole, mid periphery and far periphery) at both superior and inferior regions of the retina (B). * represents significance between STZ PTP1B fl/fl vs STZ LysM-PTP1B group, p < 0.05. Data are expressed as mean ± standard deviation (SD) (n = 4 to 7 mice per group). One-way ANOVA analysis was performed to indicate the statistical significance followed by Tukey’s multiple comparison post-hoc test. (*P < 0.05; **P < 0.01; ***P < 0.001; ****P < 0.0001; ns, non significance). RGC, retinal ganglion cell layer; IPL, inner plexiform layer; INL, inner nuclear layer; OPL, outer plexiform layer; ONL, outer nuclear layer; PS, photoreceptor segment.

**Figure 2.**
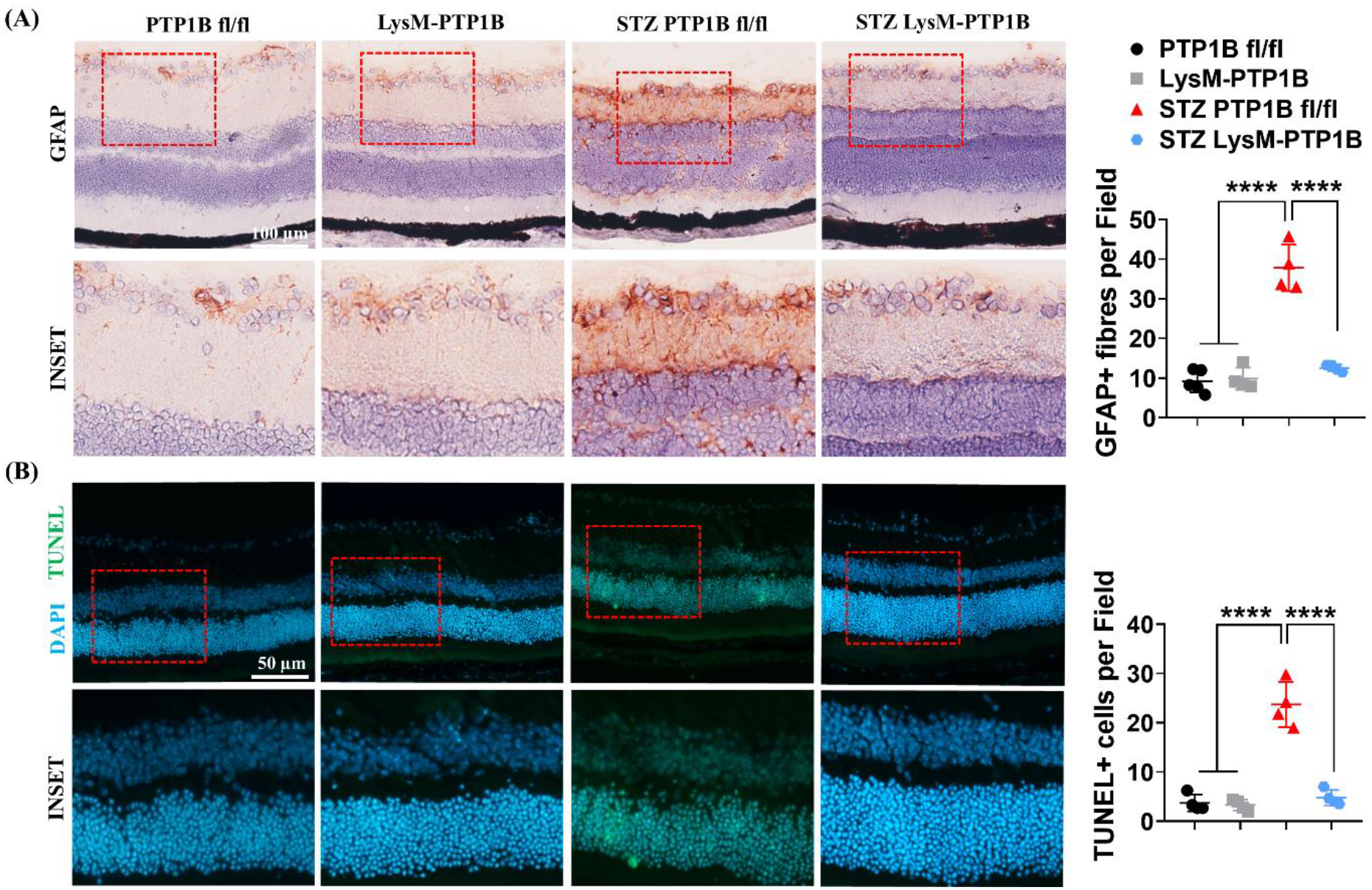
STZ-induced glial cell activation and retinal cell apoptosis is prevented in LysM-PTP1B mice. GFAP stained frozen sections from thirty-week-old male non-diabetic and diabetic LysM-PTP1B and control PTP1B-floxed mice. Scale bar: 100 µm (A). TUNEL+ cells in retinal sections of thirty-week-old male non-diabetic and diabetic LysM-PTP1B and control PTP1B-floxed mice (B). Scale bar: 50 µm. Data are expressed as mean ± SD (n = 4 to 5 mice per group). Statistical significance determined by one-way ANOVA with Tukey’s multiple comparison post-hoc test. (*P < 0.05; **P < 0.01; ***P < 0.001; ****P < 0.0001).

Deletion of PTP1B in myeloid cells prevented retinal neurodegeneration in STZ LysM-PTP1B mice compared to STZ PTP1B-floxed mice. Retinal thickness levels approached normal levels particularly towards the posterior pole of the eye, while there was less protective effect on retinal structure seen in the periphery, especially in the inferior retina in females (Fig. 1B). Upregulation of GFAP in glial cell fibres was markedly decreased in STZ LysM-PTP1B mice (Fig. 2A; Supplementary Fig. 3A) as was the number of TUNEL positive cells in the retina (Fig. 2B; Supplementary Fig. 3B). In effect, GFAP and TUNEL expression were restored to normal levels in STZ LysM-PTP1B mice in both male and female mice.

Interestingly, there was no correlation with fasted blood glucose (Supplementary Fig. 4). or other metabolic parameters such as body weight, fat and lean body mass. Both male and female STZ PTP1B-floxed mice became diabetic by 2-3 weeks post STZ-injection, while myeloid cell PTP1B-depleted mice, if anything, became diabetic slightly earlier. This was accompanied with changes in body weight and mass which recovered slowly in female mice but less so in male mice (Supplementary Fig. 5)

### Microglial cell activation and development of acellular capillaries in STZ-induced DR is regulated by myeloid cell PTP1B

STZ-induced DR is associated with inflammatory signs in the retina (38) including activation of microglia. Microglia are yolk-sac derived, self-renewing resident myeloid cells in the brain and retina, and are considered to be tissue re-modelling and waste-disposal cells. In STZ-induced DR, they become activated, evidenced by a change in morphology from a ramified cell with numerous prominent processes to a rounded, phagosome-rich cell with few processes. We have evaluated and quantified these morphological changes in microglia using an automated image analysis programme (see Methods) and confirmed that while in STZ PTP1B-floxed mice the number of Iba-1+ microglia remains unchanged compared to control mice, the number of processes (endpoints) per cell as well as the length of the processes is markedly reduced, indicating substantial microglial activation. These effects were greatly decreased in STZ LysM-PTP1B mice and were similar in both male and female mice (Fig. 3A and B; Supplementary Fig. 6A and B).

**Figure 3.**
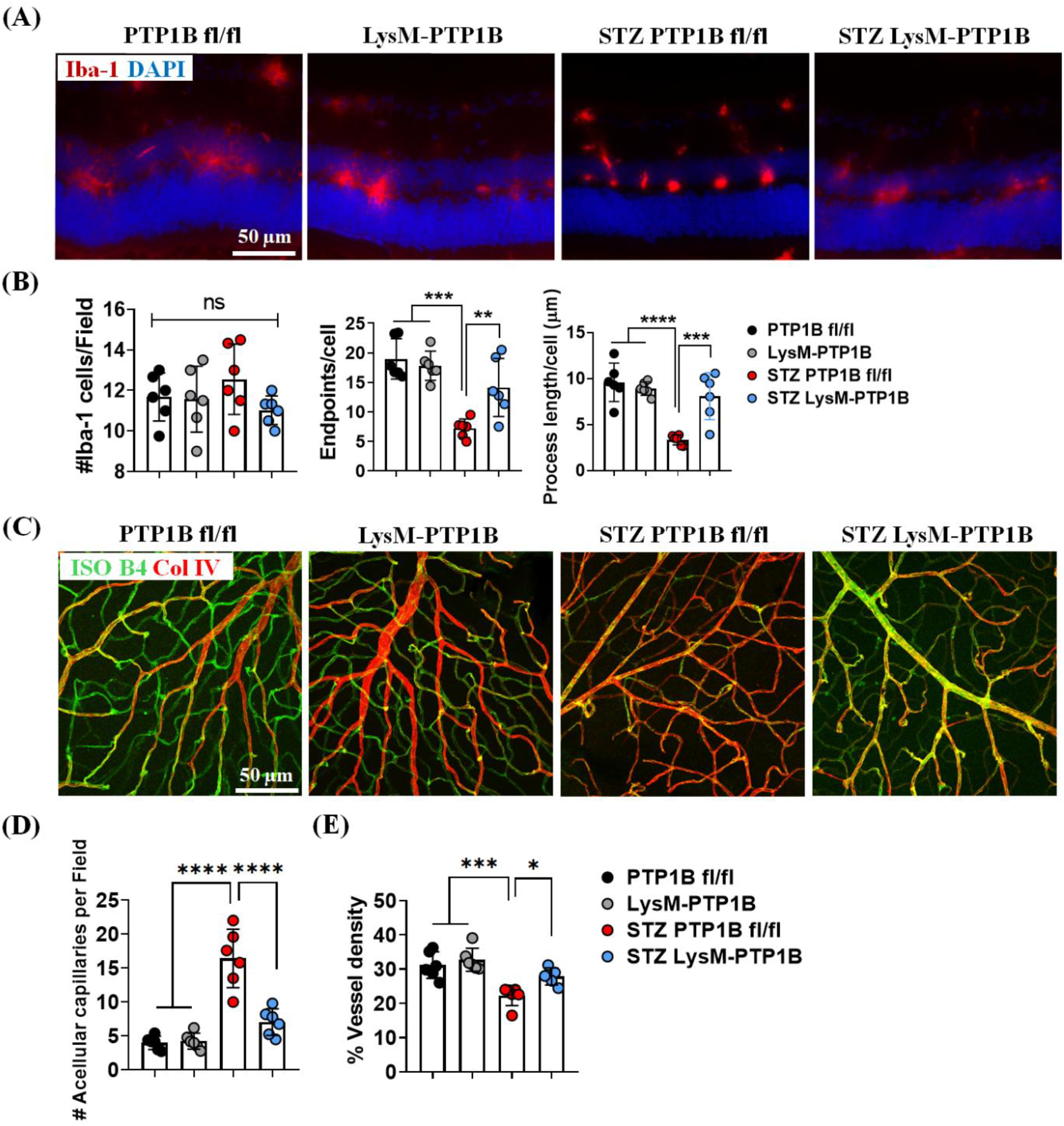
LysM-PTP1B regulates retinal microglial cell activation and development of acellular capillaries in STZ mice. Representative images of frozen sectioned retinas from thirty-week-old male non-diabetic and diabetic LysM-PTP1B and control littermate PTP1B-floxed mice using Iba-1 staining (A). Note the reduction in microglial processes in STZ mice and normal ramified appearance of microglia in LysM-PTP1B mice. Scale bar: 50 µm. Quantitative data on microglial morphology: number of Iba-1+ cells per X20 field of view, number of processes (endpoints) and process length (indicator of ramification) per cell in non-diabetic and diabetic retinas of thirty-week-old male mice (B). Representative images of retinal flatmount stained with anti-isolectin B4 (green) and anti-collagen IV (red) in thirty-week-old male non-diabetic and diabetic LysM-PTP1B and control littermate PTP1B-floxed mice (C). Quantification of acellular capillaries (D) and vessel density (E) in flatmounts of thirty-week-old male non-diabetic and diabetic LysM-PTP1B and control littermate PTP1B-floxed mice. Scale bar: 50 µm. Data are represented as mean ± SD (n = 5 to 6 mice per group). Statistical significance assessed by one-way ANOVA with Tukey’s multiple comparison post-hoc test. (*P < 0.05; **P < 0.01; ***P < 0.001; ****P < 0.0001).

Loss of endothelial cells and acellular capillaries are considered early events in the vasculopathy in DR (39,40). These were evaluated by immunostaining retinal flatmounts with antibody against collagen IV in vascular basement membrane and isolectin B4, a marker for endothelium. Loss of vascular endothelium in acellular capillaries can be detected in vessels which stain positive for collagen IV and negative for isolectin B4. The number of acellular capillaries was increased in STZ PTP1B-floxed mice compared to non-diabetic mice but greatly decreased in STZ LysM-PTP1B mice (Fig. 3C and D; Supplementary Fig. 7A and B). Similar results were obtained for overall vessel density indicating that myeloid cell PTP1B-depletion prevented endothelial cell loss and restored vessel integrity in retinas from diabetic mice (Fig. 3E; Supplementary Fig. 7C).

### Systemic inflammation in STZ-induced diabetes is decreased in myeloid cell PTP1B-depleted mice

We have previously suggested that circulating activated CCR5+CD11b+ myeloid cells were the agents of damage in peripheral tissues such as the retina, as part of a systemic inflammatory response to STZ in which bone marrow-derived myeloid cells are activated *in situ*, egress to the circulation and migrate through / populate the tissues (8). In this study, we compared inflammatory cell markers by flow cytometry in BM vs spleen cells using spleen as a surrogate model of a peripheral tissue. We gated on CD45+CD11b+ cells and initially assessed expression of activation markers, namely CD86 and MHC Class II (Supplementary Fig. 8).

Induction of diabetes using STZ was associated with increased expression of CD86 and less so MHC Class II on CD45+CD11b+ cells in both BM and spleen in male and female mice (Fig. 4A and C; Supplementary Fig. 9A and C). Deletion of myeloid cell PTP1B generally led to a decrease in the number of CD45+CD11b+ activated cells. Similar findings were observed in male mice with regards to chemokine receptor expression (CCR2, CCR5, CX3CR1) (Fig. 4B and D) but in female mice there was no difference in CX3CR1 expression in LysM-PTP1B vs PTP1B-floxed mice with and without diabetes (Supplementary Fig. 9B and D).

**Figure 4.**
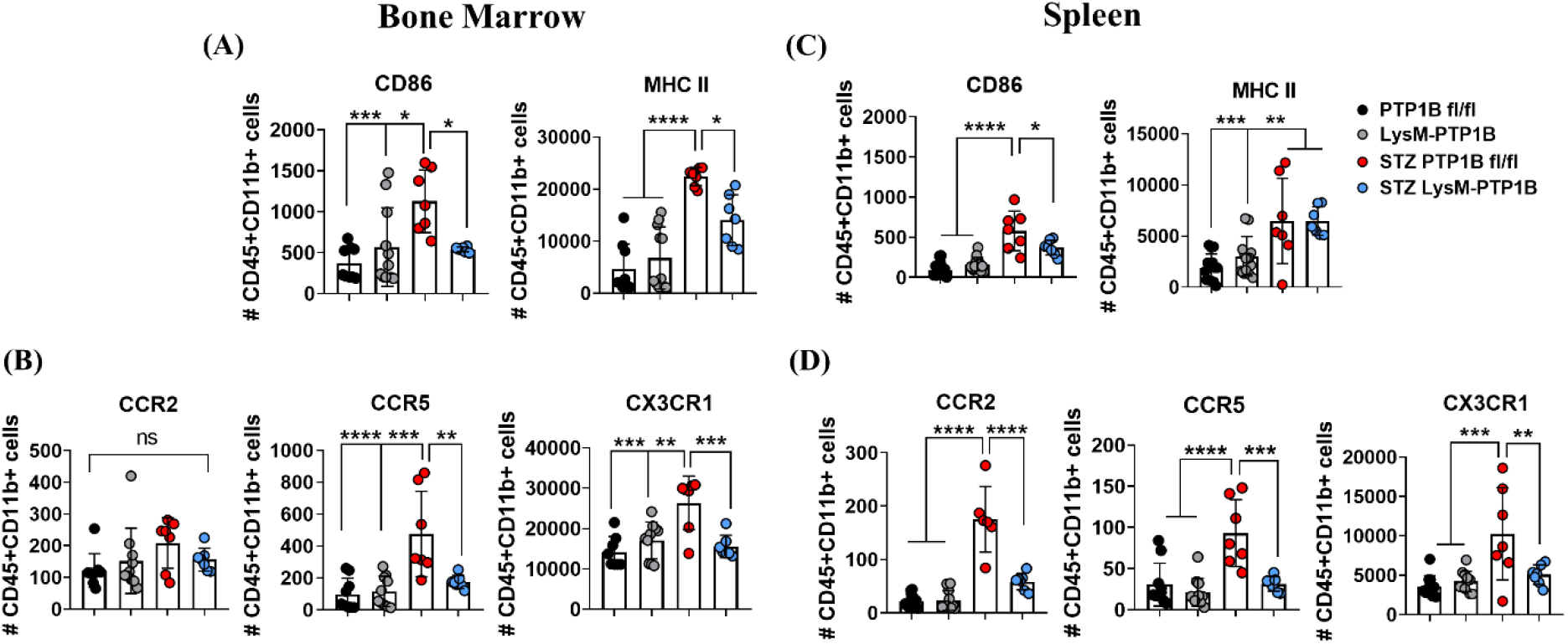
STZ LysM-PTP1B mice express lower levels of myeloid cell activation. Histogram scatter plots of absolute number of CD86 and MHC II expressing on CD45+CD11b+ cells from thirty-week-old male mouse BM (A) and spleen (C). STZ-induced increased levels of CD86 and MHC Class II are decreased in LysM-PTP1B mice. Histogram scatter plots denote the absolute number of chemokine receptors show similar decrease in chemokine receptor expression (B and D). Data are mean ± SD (n = 5 to 19 mice per group). Statistical significance assessed by one-way ANOVA with Tukey’s multiple comparison post-hoc test. (*P < 0.05; **P < 0.01; ***P < 0.001; ****P < 0.0001).

### The small molecule PTP1B inhibitor, MSI-1436 prevents STZ-induced retinal degeneration

The above data demonstrated that myeloid cell PTP1B-deletion prevented retinal neurodegeneration, as well as canonical signs of diabetic retinopathy (capillary endothelial cell loss). We wished therefore to see whether broad spectrum pharmacological inhibition of PTP1B would have similar effects. Mice were rendered diabetic with STZ as described above and five weeks post diabetes were commenced on a course of weekly injections of MSI-1436 (1 mg / kg) for eight weeks when the mice were killed (Supplementary Fig. 10). The STZ-induced loss of retinal thickness, increased GFAP expression and increased TUNEL staining were all decreased by MSI-1436 (Trodusquemine) treatment (Fig. 5A-C) as was microglial cell activation (Fig. 6A and B). Interestingly, the small molecule PTP1B inhibitor, MSI-1436 at this dose had no effect on STZ-induced changes in metabolic parameters such as fasted blood glucose, body weight, fat or lean mass (Supplementary Fig. 11) similar to the findings in STZ LysM-PTP1B mice (Supplementary Fig. 4 and 5).

**Figure 5.**
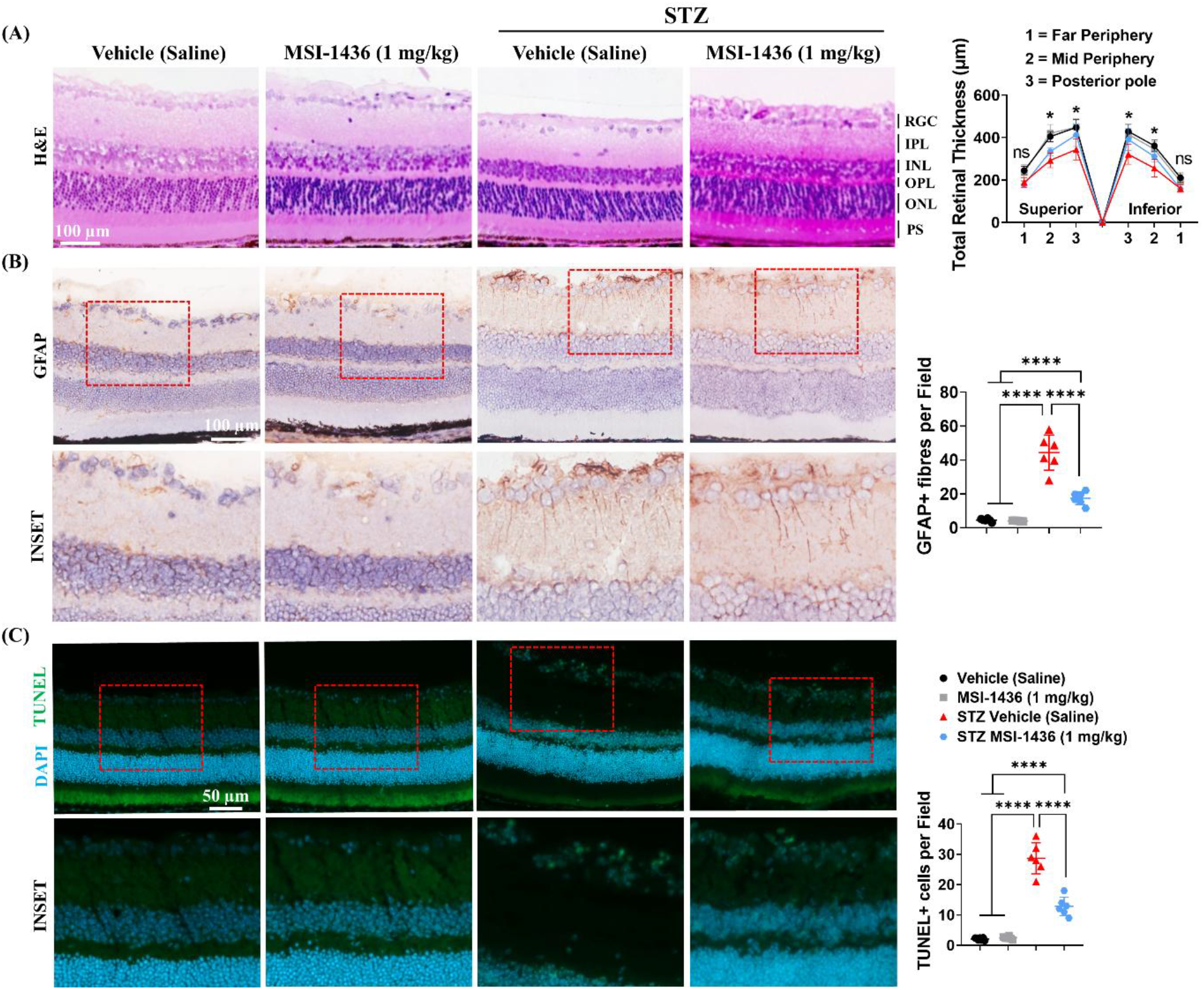
The small molecule PTP1B inhibitor, MSI-1436 rescued the retinal neurodegeneration in STZ mice. H&E, GFAP, and TUNEL staining of frozen retinal sections to assess retinal thickness (A), glial cell activation (B), and apoptotic cell death (C) of C57BL/6J mice treated with saline or MSI-1436 in physiological and diabetic conditions. In retinal thickness plots, * indicates significance between STZ Vehicle (saline) vs STZ MSI-1436 (1 mg/kg), p < 0.05. Scale bar indicates 100 µm for HE and GFAP staining, while 50 µm for TUNEL staining. Values are expressed as mean ± SD (n = 6 to 9 mice per group). Statistical significance analysed by one-way ANOVA with Tukey’s multiple comparison post-hoc test. (*P < 0.05; **P < 0.01; ***P < 0.001; ****P < 0.0001).

**Figure 6.**
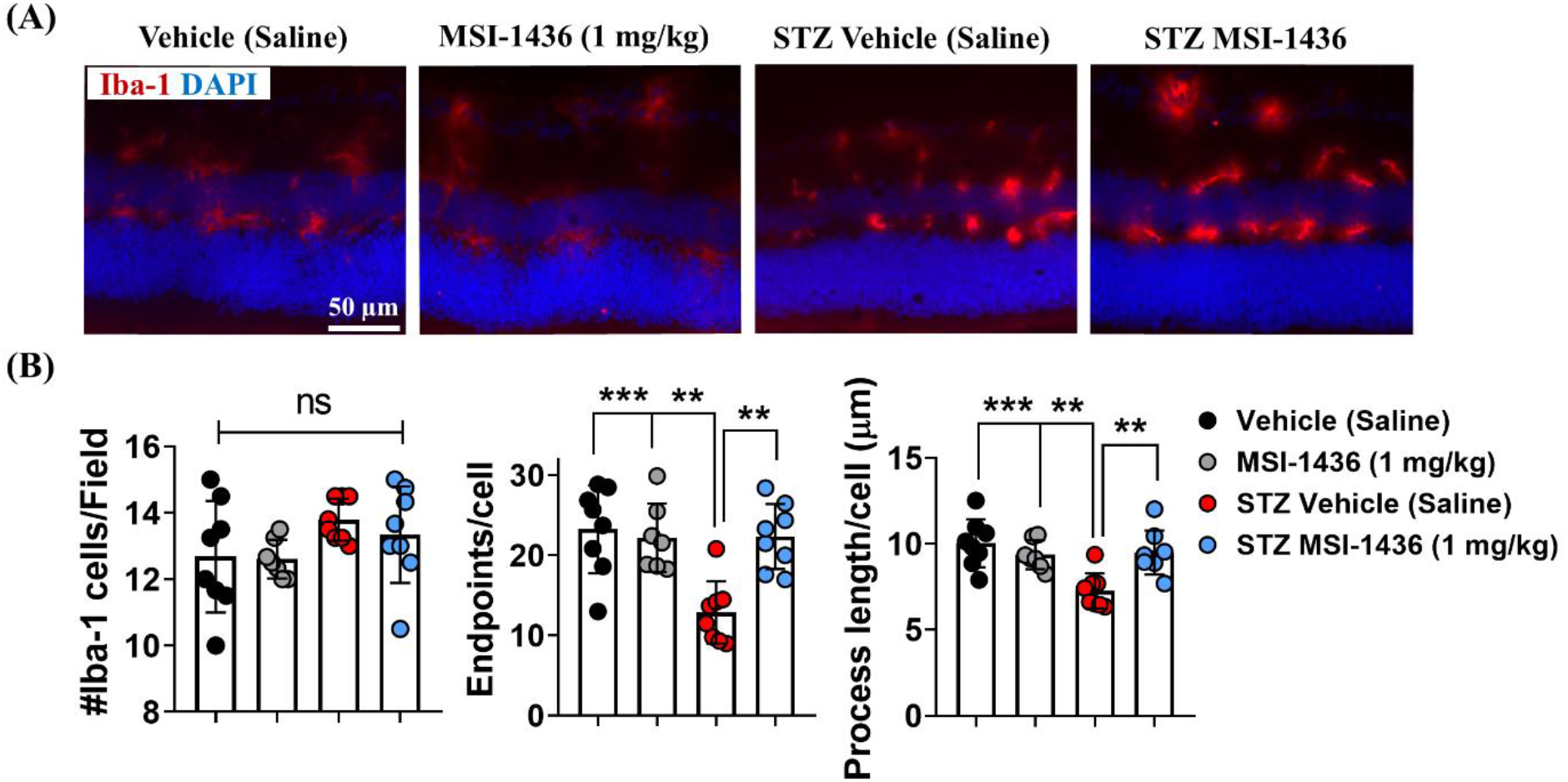
The small molecule PTP1B inhibitor, MSI-1436, decreased microglial cell activation in STZ mice. Representative images of microglia in retinas of non-diabetic and diabetic saline or MSI-1436 treated mice (A). Scale bar: 50 µm. Quantitative data on microglial morphology: number of Iba-1+ cells per X20 field of view, number of processes (endpoints) and process length (indicator of ramification) per cell in non-diabetic and diabetic retinas of male mice (B). Values are expressed as mean ± SD (n = 7 to 8 mice per group). Statistical significance analysed by one-way ANOVA with Tukey’s multiple comparison post-hoc test. (*P < 0.05; **P < 0.01; ***P < 0.001; ****P < 0.0001).

STZ-induced CD45+CD11b+ BM macrophage activation (CD86) was also decreased by MSI-1436 treatment, but not spleen macrophages (Supplementary Fig. 12A and C). Chemokine receptor expression from BM was also decreased indicating that MSI-1436 was having a direct effect on chemokine receptor induction. However, there was a limited effect of PTP1B inhibition on splenic CCR2 and CCR5, indicating that the drug’s main effect was on the phase of receptor induction in the BM (Supplementary Fig. 12B and D). Surprisingly, STZ itself had little effect on splenic activation markers (CD86 and MHC Class II).

### Deletion and inhibition of PTP1B prevents high glucose-induced macrophage apoptosis and mitochondrial dysfunction in bone marrow-derived macrophages *in vitro*

The above data demonstrate that myeloid cell PTP1B is involved in STZ-induced retinal neurodegeneration, possibly through myeloid cell activation, in particular suggesting a direct role in the induction of chemokine receptors on bone marrow myeloid cells (Fig. 3A and B; Supplementary Fig. 12B). We therefore wished to investigate possible mechanisms using cultured bone marrow-derived macrophages (BMDMs). BMDMs were isolated and cultured *in vitro* in the presence of normal (5mM) or high glucose (25mM) as described in Methods. Exposure of BMDMs to high glucose induced increased levels of apoptosis (10%), an effect which was decreased in LysM-PTP1B cells (Fig. 7A). This correlated with marked decrease in mitochondrial membrane potential and an increase in superoxide production, both of which were prevented in PTP1B-deficient macrophages (Fig. 7B and C). We also assessed the effect of the PTP1B inhibitor, MSI-1436 on high glucose challenged BMDMs. Preliminary experiments revealed that MSI-1436 was non-toxic to BMDMs in a dose range of 10-50 nM (data not shown). Treatment of BMDMs with 30 nM MSI-1436 completely prevented high glucose-induced apoptosis (Fig. 7D). In addition, both lower mitochondrial membrane potential and increase in superoxide production were prevented (Fig. 7E and F). Together, the data suggests that PTP1B actively mediates diabetes-induced retinal neurodegeneration through high glucose-induced mitochondrial dysfunction in myeloid cells and that PTP1B represents a valid therapeutic target in the early stages of DR.

**Figure 7.**
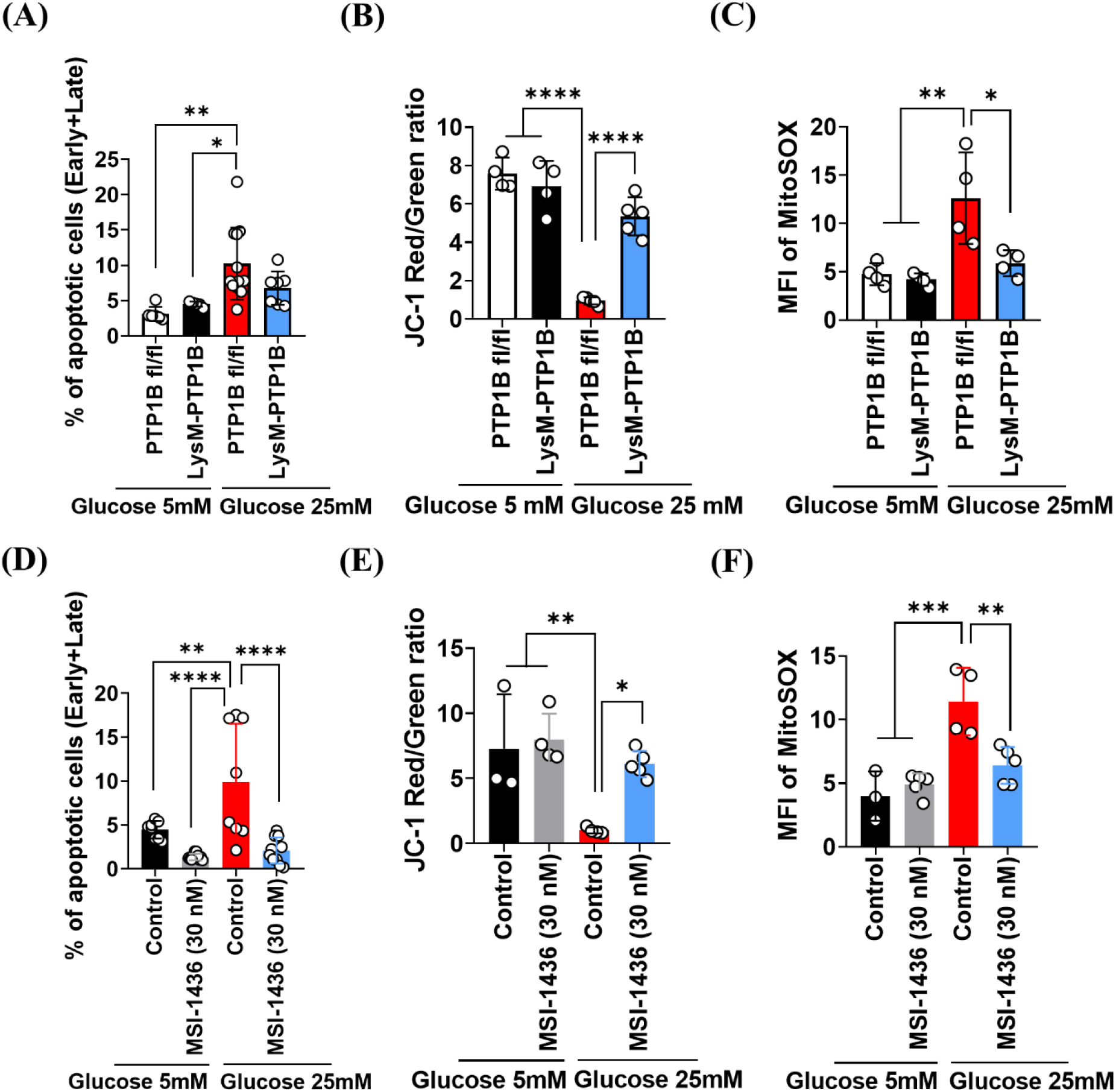
Both Deletion and Inhibition of PTP1B prevent apoptotic cell death and accumulation of dysfunctional mitochondria in BMDMs challenged with high glucose. BMDMs isolated from non-diabetic and diabetic male LysM-PTP1B and control littermate PTP1B-floxed mice seeded in normal (5 mM) and high glucose (25 mM) conditions for 7 days were harvested and incubated with annexin V and propidium iodide (PI), JC-1 and MitoSOX reagents to evaluate apoptotic cell death (A) mitochondrial membrane potential (ΔΨm) (B), and mitochondrial superoxide production (C) using flow cytometry and fluorescence microscopy. BMDMs isolated from non-diabetic and diabetic C57BL/6J mice treated with MSI-1436 (30 nM) for 24 h seeded in normal and high glucose medium were harvested and probed with PI and annexin V, JC-1, and MitoSOX reagents to assess apoptotic cell death (D), mitochondrial membrane potential (E), and mitochondrial superoxide generation (F). Scale bar: 50 µm. Data are represented as mean ± SD (n = 3 to 12 mice per group). Statistical significance analysed by one-way ANOVA with Tukey’s multiple comparison post-hoc test. (*P < 0.05; **P < 0.01; ***P < 0.001; ****P < 0.0001).

## Discussion

Diabetes mellitus, both type 1 and type 2, causes major complications (retinopathy, nephropathy, neuropathy) due to poorly regulated glucose and lipid metabolism (41). Evidence abounds that these complications are caused by chronic medium-to-low grade inflammation while recent data suggest that, at least in the case of retinopathy (DR), direct tissue damage (neurodegeneration) may be due to metabolic effects of glucose, lipid and their metabolites (42,43). The early onset of retinopathy in diabetes has been cited as evidence for a direct metabolic effect of high blood glucose on the retina, since DR is classically considered to develop as a cumulative, progressive pathology (reviewed in reference (13)). However, vascular occlusion by activated leukocytes (leukostasis) occurs within two weeks of onset of diabetes, at least in experimental models (8), and so the question remains unresolved i.e. whether retinal damage is the result of diabetes-induced inflammatory processes or is a direct metabolic complication.

In this study, we have investigated the role of inflammatory myeloid cells in the induction of DR. Specifically, we have explored the role of myeloid cell PTP1B in early onset of DR (neurodegeneration). PTP1B is recognised as a driver of diabetes where, as a regulator of the insulin receptor signalling, it controls the level of tissue glucose uptake. PTP1B is a universally expressed enzyme, and several tissue- and cell-specific PTP1B-deleted models have been described (44). Previously, we have shown that myeloid cell-specific PTP1B deletion (LysM-PTP1B) in mice leads to decreased inflammatory responses in response to lipopolysaccharide both *in vivo* and *in vitro* (28,29). In the current study we observed that while STZ-mice developed significant levels of diabetes, deletion of PTP1B was not accompanied by improvement in fasted blood glucose. Despite the lack of effect on diabetes control which represents more closely type 1 diabetes, neurodegeneration (retinal thinning) induced by STZ in control PTP1B fl/fl mice was prevented in STZ-induced diabetic LysM-PTP1B mice. Similarly, there was less Muller glial cell (GFAP) and microglial cell activation (dendrite expression), as well as fewer signs of neuronal apoptosis as determined by TUNEL staining in both, male and female mice. The protective effects of PTP1B deletion on retinopathy in the STZ LysM-PTP1B mice were also observed in STZ C7BL/6J mice treated with the small molecular PTP1B inhibitor, MSI-1436. Both retinal neurodegeneration and local glial cell activation were decreased, but there was no difference in overall glucose metabolism or in body weight in MSI-1436 treated STZ mice compared to untreated STZ mice. This has allowed us to delineate between the effects of PTP1B deletion on glycaemia and body weight versus direct regulation of inflammation and neurodegeneration.

Capillary loss (drop-out) is a signal feature of DR. In this study we observed that STZ mice lost capillary density and sustained larger numbers of acellular capillaries, as an early feature of disease. Interestingly, capillary integrity was protected in LysM-PTP1B mice (Fig. 3C-E) despite persistence of severe STZ-induced hyperglycaemia (Supplementary Fig. 4). Whether the protective effect of PTP1B deletion was due to decreased microglial activation or reduced leukostasis is not clear since both mechanisms can cause endothelial cell damage and loss of capillaries (8). The data in the present work suggests that both, activated microglia and circulating leukocytes may contribute to the diabetic vasculopathy, while the neurodegeneration is more likely to be the direct result of loss of physiological local microglial function (e.g. synaptic trimming).

These findings thus support a local effect of PTP1B on signs of retinopathy, despite the fact that there was downregulation of the STZ-induced systemic inflammatory response in both the myeloid cell PTP1B-depleted mice and the PTP1B inhibitor treated mice (Fig. 1-3, Fig. 5 and 6). Microglial cells and retinal pigment epithelium (RPE) both express LysM and PTP1B and it is likely that PTP1B is decreased in these cells in the retina. Microglia have a major physiological housekeeping role in the central nervous system particularly in synaptic trimming while RPE cells transport recycled retinal material to the choroidal vasculature (45– 47). Microglia are also activated by stress and inflammatory stimuli as seen here, whereby there is a major shift in morphology from elaborate dendritiform cells to rounded cells (Figure 3A and B).

A search for possible mechanisms of action of PTP1B on myeloid cell physiology revealed that in culture, PTP1B depleted macrophages, as well as MSI-1436 treated macrophages, were more resistant to apoptosis and loss of membrane potential, as well as lower producers of superoxide in the presence of 25 mM glucose, indicating that PTP1B is required for a positive proinflammatory response by myeloid cells.

In summary, while PTP1B is recognised as a major driver of insulin resistance (48) and Type 2 diabetes via its effect on the insulin receptor dephosphorylation, its expression in myeloid cells has little effect on glucose control under hyperglycaemic, type 1 diabetes conditions, while it mediates a pro-inflammatory, damaging effect on retinal neurodegeneration and retinal capillary vessel integrity. The evidence suggests that PTP1B has both, a local effect on microglia and a systemic effect on bone marrow-derived circulating myeloid cells. The data also suggest that, at least in this model of diabetes, the pro-inflammatory effect of PTP1B-expressing myeloid cells, rather than any metabolic hyperglycaemic effect, directly causes the retinal complications. The inhibitory effects of global inhibition of PTP1B using MSI-1436 further suggest that its mode of action may also be on signalling through inflammatory pathways and is a promising indicator for the use of such agents in disorders associated with inflammaging (49,50).

## Article information

## Acknowledgements

The authors acknowledge the University of Aberdeen Histology and Microscopy core facility, the Iain Fraser Flow Cytometry Core facility and Medical Research Facility for their assistance and guidance.

## Funding

This study was supported by funds from the BHF project grant to M.D., J.V.F and L.K. (PG/21/10555).

## Duality of Interest

Authors declare no conflict of interest

## Author Contributions

Conceptualization: A.A., L.K., J.V.F., M.D.; Methodology: A.A., M.B., A.O., S.K-S.; Validation and Formal analyses: A.A., M.B., A.O., S.K-S., L.K., J.V.F., M.D.; Investigation: A.A.; Resources and Data curation: A.A., L.K., J.V.F., M.D.; Writing-original draft and Writing-review and editing: A.A., L.K., J.V.F., M.D.; Visualization A.A.; Supervision, Project administration and Funding acquisition: L.K., J.V.F., M.D.; M.D. is the guarantor of this work and, as such, had full access to all the data in the study and takes responsibility for the integrity of the data and the accuracy of the data analysis.

## Notes

### Competing Interest Statement

The authors have declared no competing interest.

