## Supplementary material for "Myeloid cell protein tyrosine phosphatase 1B (PTP1B) drives retinal neurodegeneration in diabetic mice"

**Running Title: Myeloid PTP1B drives retinal neurodegeneration**

Ayaz Ali[1,2], Morgan Boyne[1,2], Abrar Othman[1,3], Sarah Kamli-Salino[1], Lucia Kuffova[2,4], John V. Forrester[2*] and Mirela Delibegovic[1*]

[1]Aberdeen Cardiovascular and Diabetes Centre, Institute of Medical Sciences, University of Aberdeen, Aberdeen, UK.

[2]Aberdeen Ophthalmology Group, Institute of Medical Sciences, University of Aberdeen, Aberdeen, UK.

[3]Department of Clinical Laboratory Sciences, Faculty of Applied Medical Sciences, Taibah University, Madinah, Saudi Arabia

[4]INSiGHT Comprehensive Eye Care, Hamilton, Bermuda

### Supplementary data

#### Mice and mouse husbandry

DNA extraction and genotyping was performed for the new litters before including them in the study. PTP1B fl/fl and mice expressing Cre under the control of the LysM promoter ($LysM^{cre}$) were maintained at the Medical Research Facility, University of Aberdeen, Aberdeen, UK. Mice were crossed to produce $LysM^{cre}$-PTP1B fl/fl (LysM-PTP1B) lacking PTP1B in myeloid cells. The mice were kept under the 12h light-dark cycle having free access to food and water. Two days before the streptozotocin (STZ) administration, mice were housed in individually ventilated cages.

#### Induction of STZ-induced diabetes model and study design

Fourteen- and twenty-two-week-old male and female mice were acclimatised to handling for two weeks and then allocated to four groups: non-diabetic PTP1B fl/fl, non-diabetic LysM-PTP1B, STZ-induced diabetic PTP1B fl/fl, and STZ-induced diabetic LysM-PTP1B. STZ was dissolved in 0.05M citrate buffer (sodium citrate dihydrate and citric acid monohydrate, pH 4.5) administered for five consecutive days via the intraperitoneal cavity. Age and gender matched non-STZ control littermates were administered with the same amount of citrate buffer. In the following week, fasted blood glucose was measured using a glucometer (AlphaTRAK, UK), and mice attaining a blood glucose level of >18 mmol/L were considered diabetic. All animal groups were monitored for body weight and fasted blood glucose weekly throughout the study. Thereafter, mice were sacrificed using $CO_2$ inhalation and cervical dislocation. Whole blood and organs were collected for further analysis.

**Body composition measurement**

Mice were placed in a plastic tube inserted into the scanner (Echo, Medical Systems, Houston, USA) and 3 consecutive independent scans of body composition per mouse were taken at 0, 3, and 6 weeks and the average value calculated.

**Flow cytometry of spleen and bone marrow (BM)**

The panel used in this study included CD45-APC (30-F11), CD11b-BUV 395 (M1/70), CCR2-APC/Fire (SA203G11), CCR5-PerCp-eF710 (HM-CCR5(7A4)), CX3CR1-PE (SA011F11), CD86-Apc/fire (GL-1), and MHC II-FITC (M5/114.15.2). Populations of interest (total leukocytes/cells) were gated on a dot plot with logarithmic side scatter vs forward scatter and subsequently cell aggregates were excluded by implementing a forward scatter pulse gate (FSC-H vs FSC-A). All analyses performed were based on live populations of cells, where unstained live cells were employed as a gating control.

**Tissue harvesting and processing**

One enucleated eye was fixed in glutaraldehyde 2.5% in PBS for histological evaluation and the other eye was snap frozen in a mould with optimal cutting temperature (OCT) medium (Fisher Scientific, UK) in isopentane on dry ice and cryosectioned into 10 μm thick slices. Retinal sections mounted on Polysine® glass slides (Thermo Fisher, UK) were air dried overnight and stored at -20 ℃ for further experiments.

**Immunohistochemistry staining of glial fibrillary acidic protein (GFAP)**

Defrosted sections fixed in cold acetone for 15 min were rehydrated in Tris buffer saline ((TBS): 90% 1 M saline, 10% $dH_2O$, 0.5% pH 6.8 tris in $dH_2O$, 0.1% Tween-20) and blocked with streptavidin and biotin blocking reagent (Vector Laboratories, 2BScientific, UK) respectively for 15 min. Retinal sections were blocked with 5% normal rabbit serum to block non-specific binding for 30 min and incubated with the primary antibody (anti-GFAP; Agilent, Dako, UK) at 1:1000 dilution for 1 h at room temperature (RT). Thereafter, tissues probed with the secondary antibody (Agilent, Dako, UK) diluted at 1:300 for 1 h were washed in TBS-T and subsequently incubated in 3% hydrogen peroxide to block the intracellular peroxidase activity for 20 min followed by streptavidin-HRP (horseradish peroxidase; Agilent, Dako, UK) treatment for 30 min at RT. Then, tissues were incubated with the Vector NovaRed substrate (Vector Laboratories, 2BScientific, UK) for signal development following the manufacturer's guidelines and counterstained using hematoxylin (Vector Laboratories, 2BScientific, UK). Finally, tissues rinsed under the tap water were dehydrated using gradient alcohol series and histoclear (National Diagnostics, UK), mounted with the mounting medium, and photomicrographed using Zeiss Axioscope5 upright microscope (Carl Zeiss AG, Oberkochen, DE).

**Terminal deoxynucleotidyl transferase dUTP nick end labelling (TUNEL) assay**

Defrosted tissue sections were fixed in 4% paraformaldehyde for 15 min at RT. Thereafter, sections were washed and probed with the TUNEL reagent (Roche Diagnostic, Penzberg, Germany) for 1h at 37°C. Next, tissues were stained with DAPI to stain the nuclei followed by washing and mounted using aqueous fluorescent mounting medium (Sigma-Aldrich, US). Finally, apoptotic cell death was measured by detecting the TUNEL+ stained cells with wavelength in the range of 450-500 nm (excitation) and 515-565 nm (emission) using fluorescence microscopy (Zeiss Imager M2 upright microscope; Carl Zeiss AG, Oberkochen, DE). The number of TUNEL+ cells were counted per X20 field of view and used for statistical analysis.

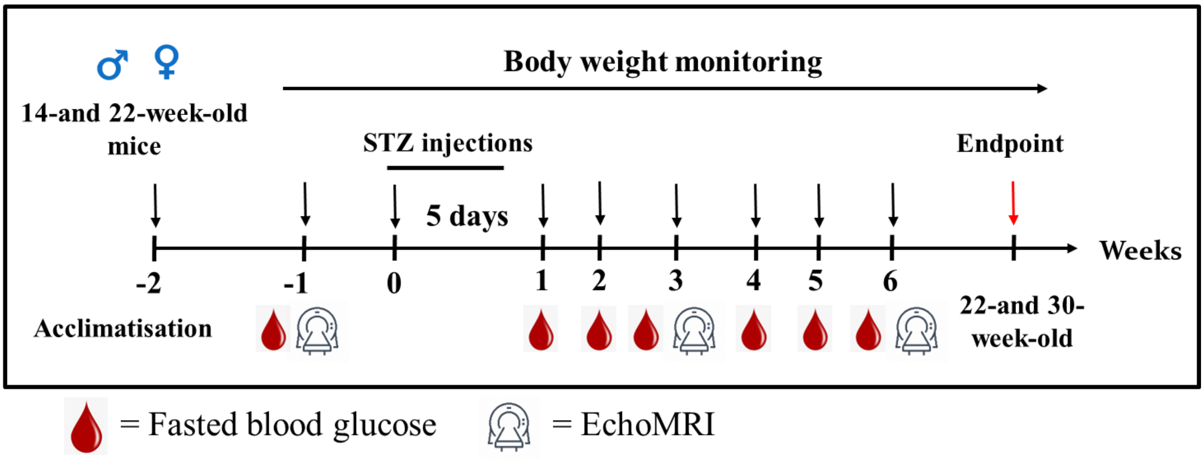


**Supplementary Figure 1. Experimental Design.** Schematic representation illustrating the timeline for induction of diabetes in fourteen- and twenty-two-week-old male and female LysM-PTP1B and control littermate PTP1B-floxed mice for six weeks. Body weight, fasted blood glucose, and body composition were assessed at designated times.

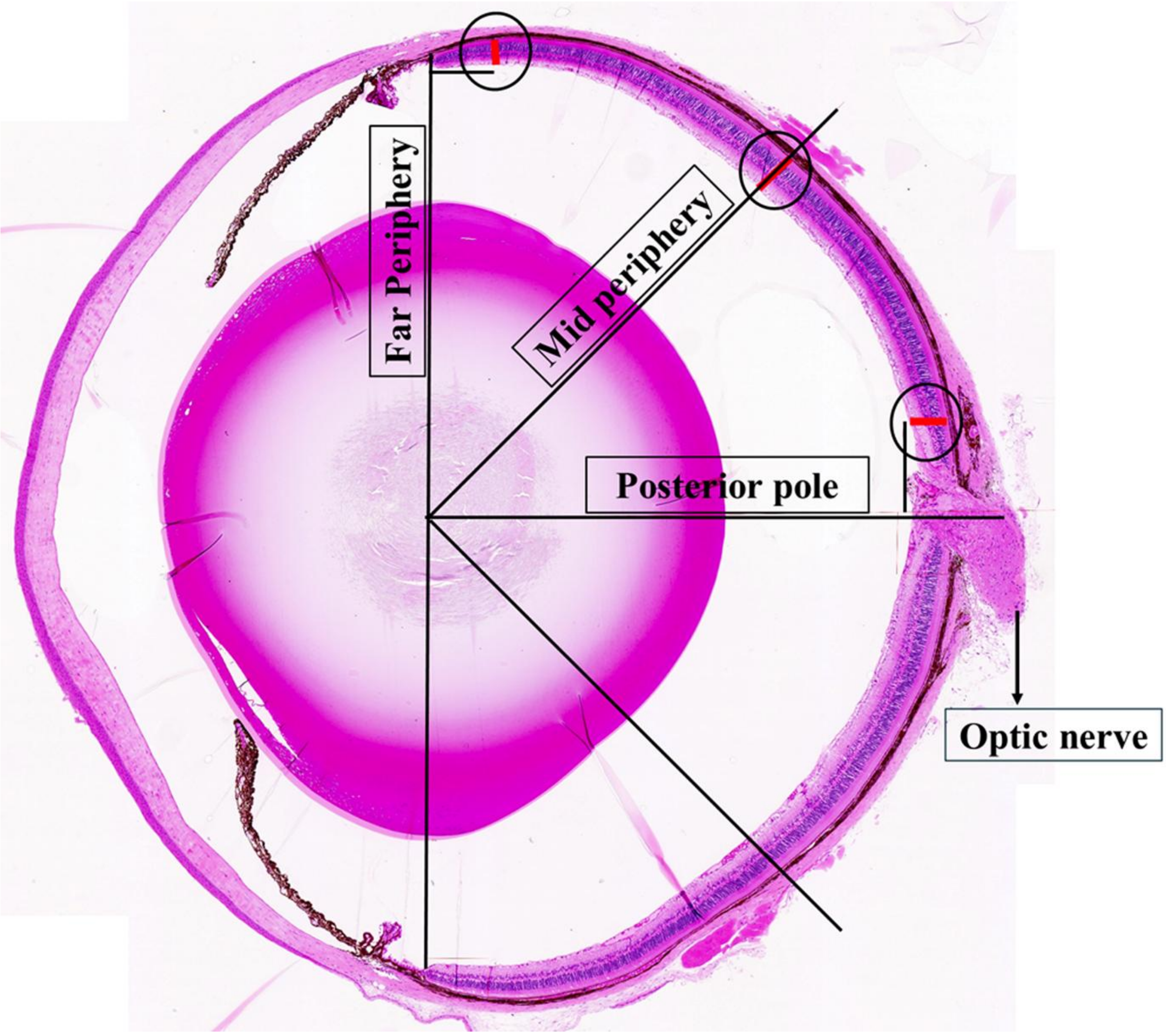


**Supplementary Figure 2. Retinal thickness measurement.** Retinal thickness was measured in resin embedded retinal sections stained with H&E at three different regions: posterior pole, mid periphery and far periphery (identified by circles).

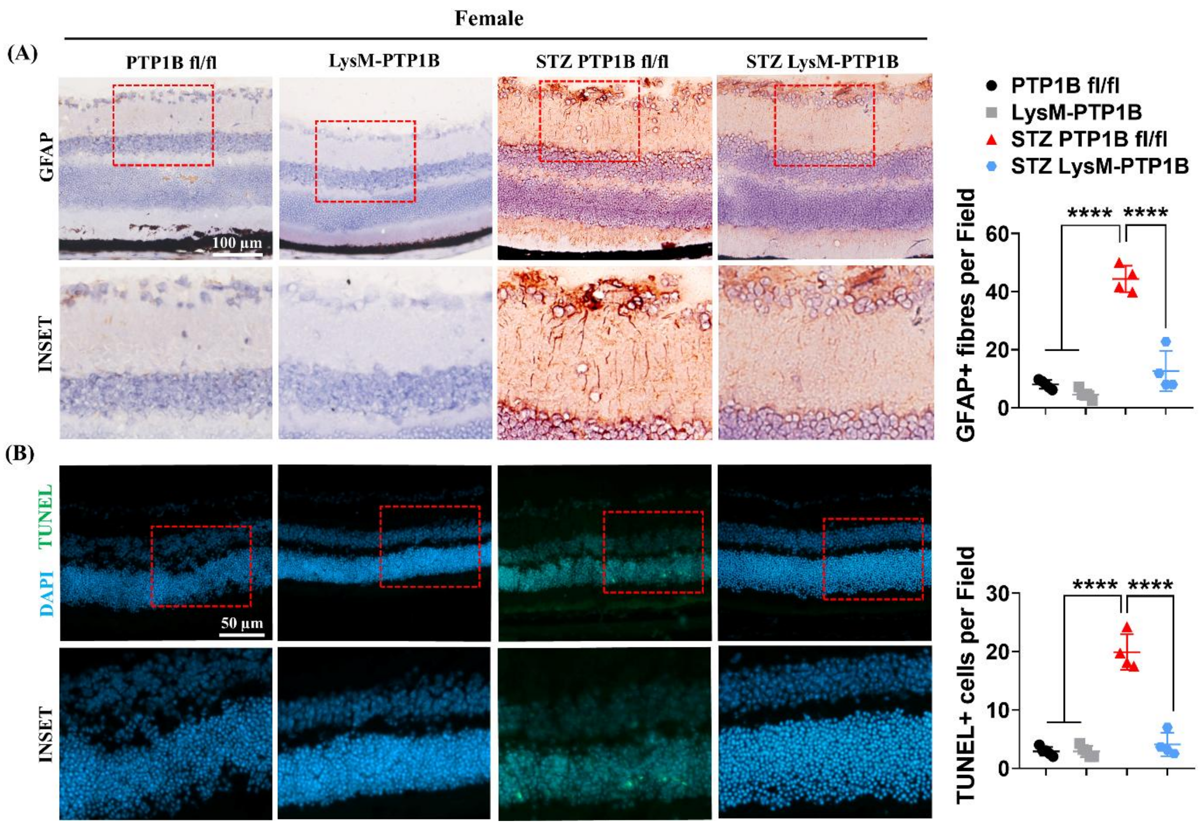


**Supplementary Figure 3. STZ-induced glial cell activation and retinal cell apoptosis is prevented in LysM-PTP1B mice.** GFAP stained frozen sections in thirty-week-old female non-diabetic and diabetic LysM-PTP1B and control PTP1B-floxed mice (A). Scale bar: 100 µm. TUNEL+ cells in retinal sections of non-diabetic and diabetic LysM-PTP1B and control PTP1B-floxed mice (B). Data are expressed as mean ± SD (n = 4 to 5 mice per group). Statistical significance determined by one-way ANOVA with Tukey's multiple comparison post-hoc test. (*P < 0.05; **P < 0.01; ***P < 0.001; ****P < 0.0001).

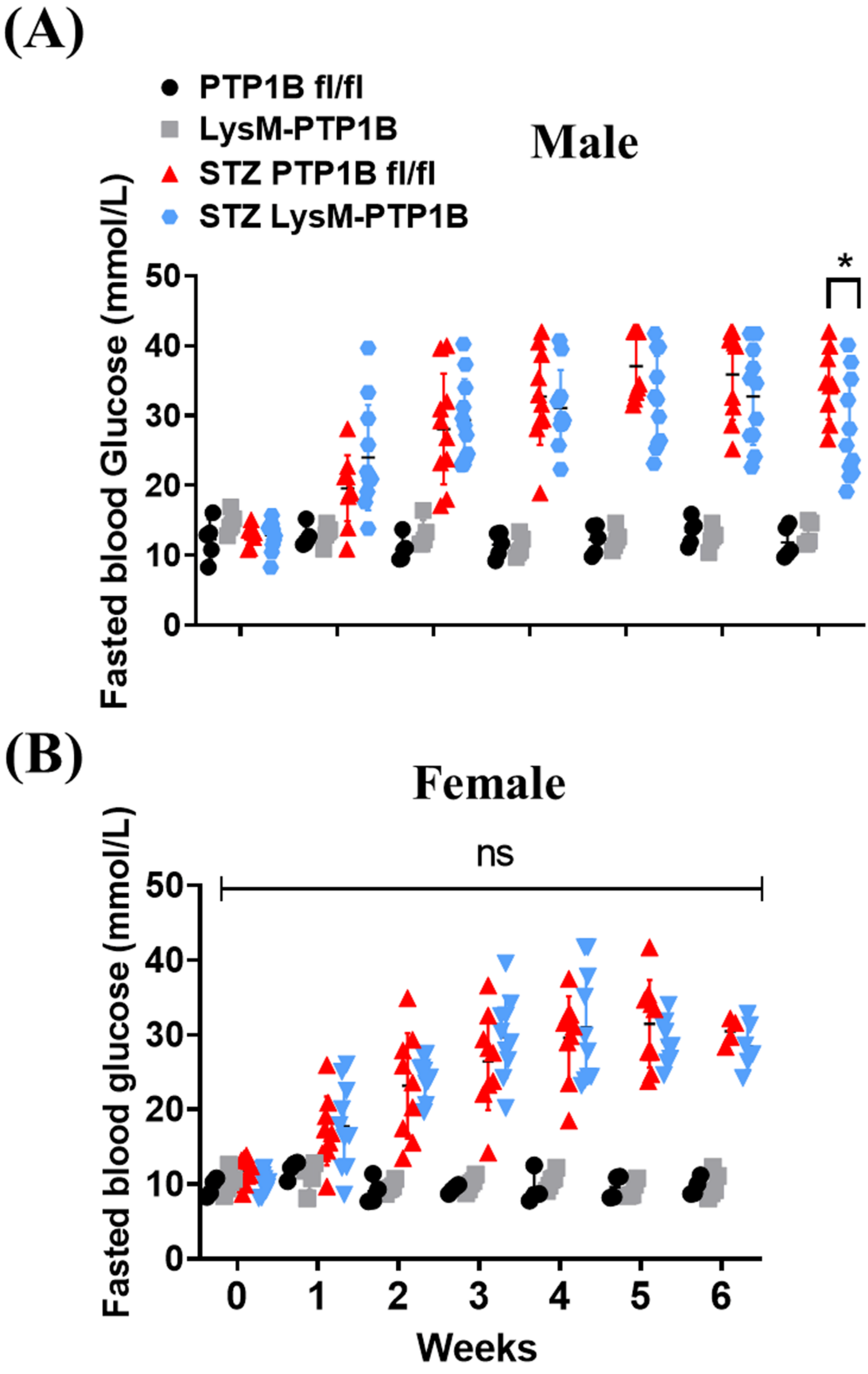
(A)
PTP1B fl/fl
LysM-PTP1B
STZ PTP1B fl/fl
STZ LysM-PTP1B
Male
Fasted blood Glucose (mmol/L)
*
(B)
Female
Fasted blood glucose (mmol/L)
ns
0
1
2
3
4
5
6
Weeks

**Supplementary Figure 4. LysM-PTP1B mice have unaltered diabetogenic effect of STZ.** Fasted blood glucose was measured in thirty-week-old male (A) and female (B) non-diabetic and diabetic LysM-PTP1B and PTP1B-floxed mice. Data are shown as mean ± SD (n = 4 to 11 mice per group). Statistical significance was assessed using two-way ANOVA followed by Bonferroni's multiple comparison post-hoc test. Black asterisks denote significance between STZ PTP1B fl/fl and STZ LysM-PTP1B group. (*P < 0.05; **P < 0.01; ***P < 0.001; ****P < 0.0001).

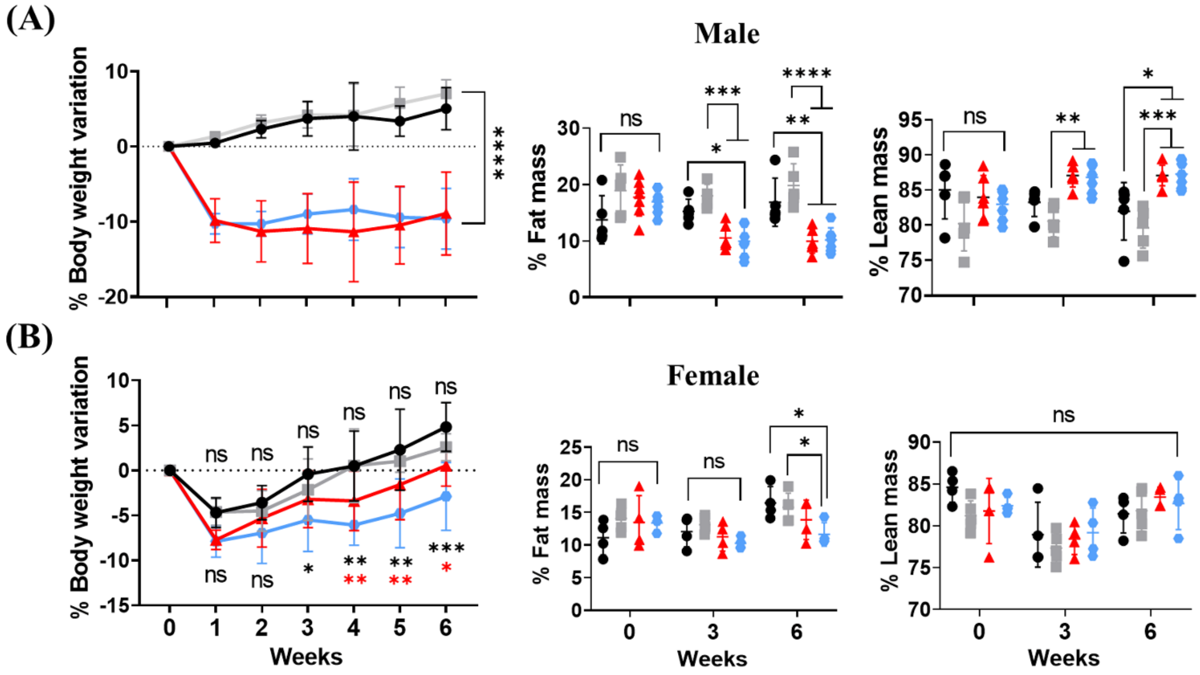


**Supplementary Figure 5. Effect of LysM-PTP1B on body weight, fat and lean mass**. LysM-PTP1B have unaltered effect on changes in body weight, fat or lean mass induced by STZ. Males (A) females (B). Data are shown as mean ± SD (n = 4 to 7 mice per group). Black asterisks represent significance compared to control PTP1B-floxed group; red asterisks indicate significance compared to control LysM-PTP1B group. Statistical significance was assessed using two-way ANOVA followed by Bonferroni's multiple comparison post-hoc test. (*P < 0.05; **P < 0.01; ***P < 0.001; ****P < 0.0001).

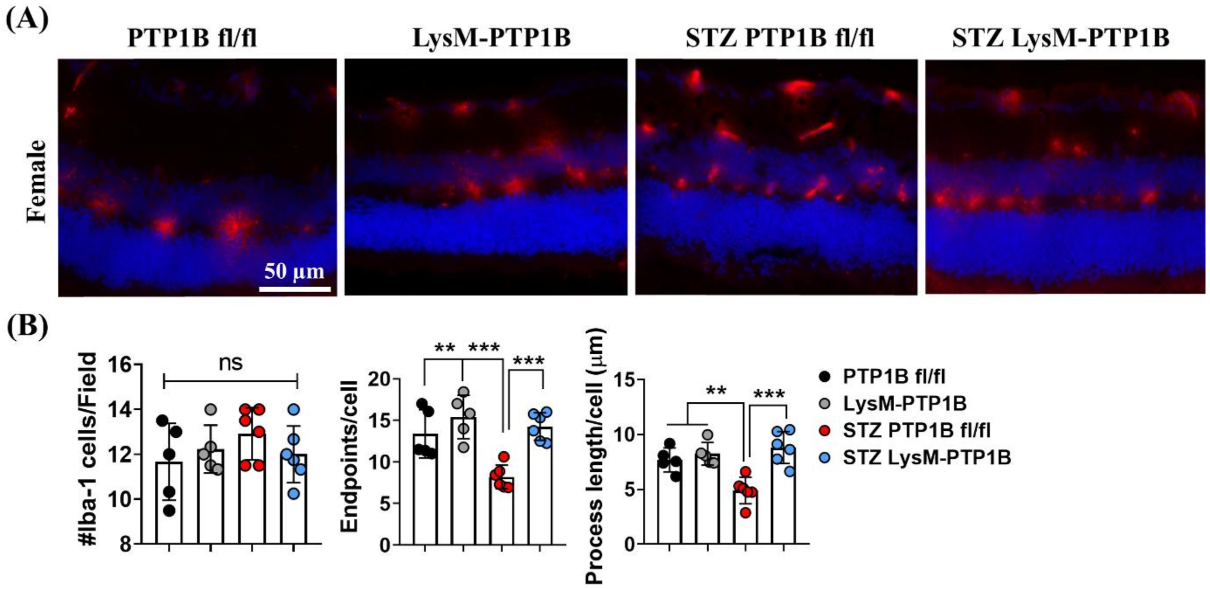


**Supplementary Figure 6. LysM-PTP1B regulates STZ-induced retinal microglial cell activation.** Representative images of frozen sectioned retinas from thirty-week-old female non-diabetic and diabetic LysM-PTP1B and control littermate PTP1B-floxed mice using Iba-1 staining (A). Note the reduction in microglial processes in STZ mice and normal ramified appearance of microglial in LysM-PTP1B mice. Quantitative data on microglial morphology: number of Iba-1+ cells per X20 field of view, number of processes (endpoints) and process length (indicator of ramification) per cell in non-diabetic and diabetic retinas of thirty-week-old female mice (B). Data are represented as mean ± SD. (n = 5 to 6 mice per group). One-way ANOVA was performed to indicate the statistical significance followed by Tukey's multiple comparison post-hoc test. (*P < 0.05; **P < 0.01; ***P < 0.001; ****P < 0.0001).

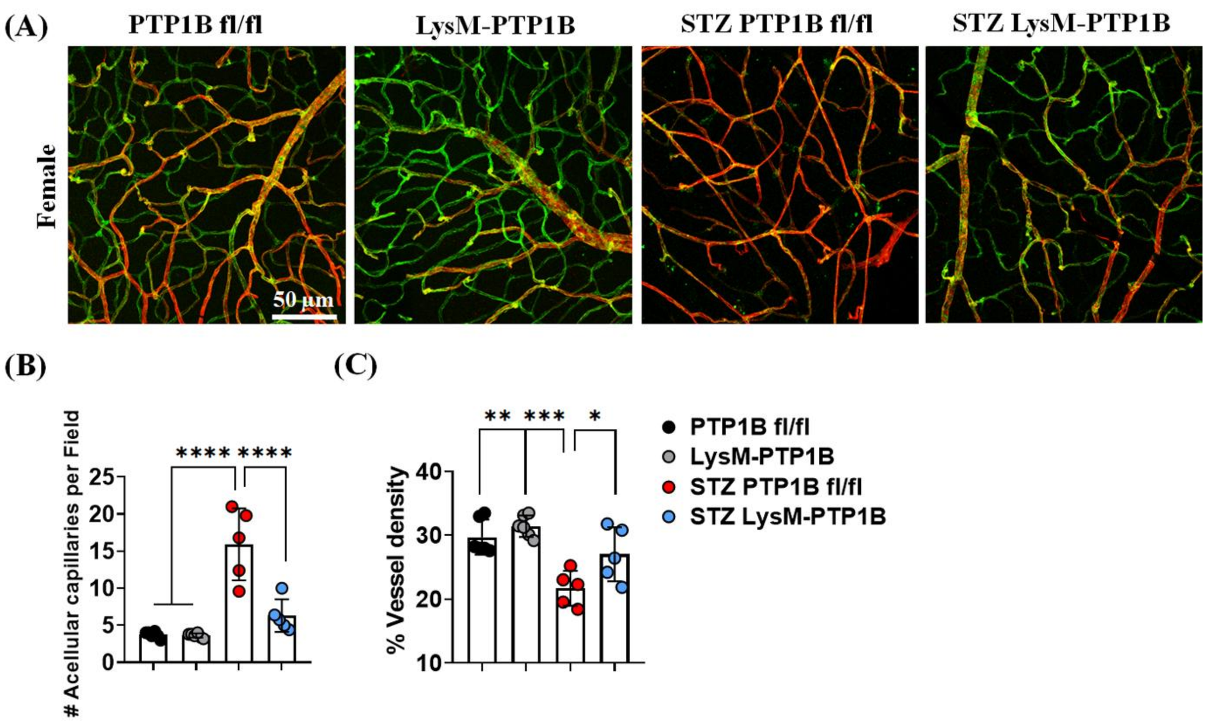


**Supplementary Figure 7. LysM-PTP1B prevents the development of acellular capillaries in STZ mice.** Representative images of retinal flatmount stained with anti-isolectin B4 (green) and anti-collagen IV (red) in thirty-week-old female non-diabetic and diabetic LysM-PTP1B and control littermate PTP1B floxed mice (A). Quantification of acellular capillaries (B) and vessel density (C) in flatmounts of thirty-week-old female non-diabetic and diabetic LysM-PTP1B and control littermate PTP1B-floxed mice. Scale bar: 50 µm. Data are represented as mean ± SD. (n = 5 to 6 mice per group). Statistical significance determined by one-way ANOVA with Tukey's multiple comparison post-hoc test. (*P < 0.05; **P < 0.01; ***P < 0.001; ****P < 0.0001).

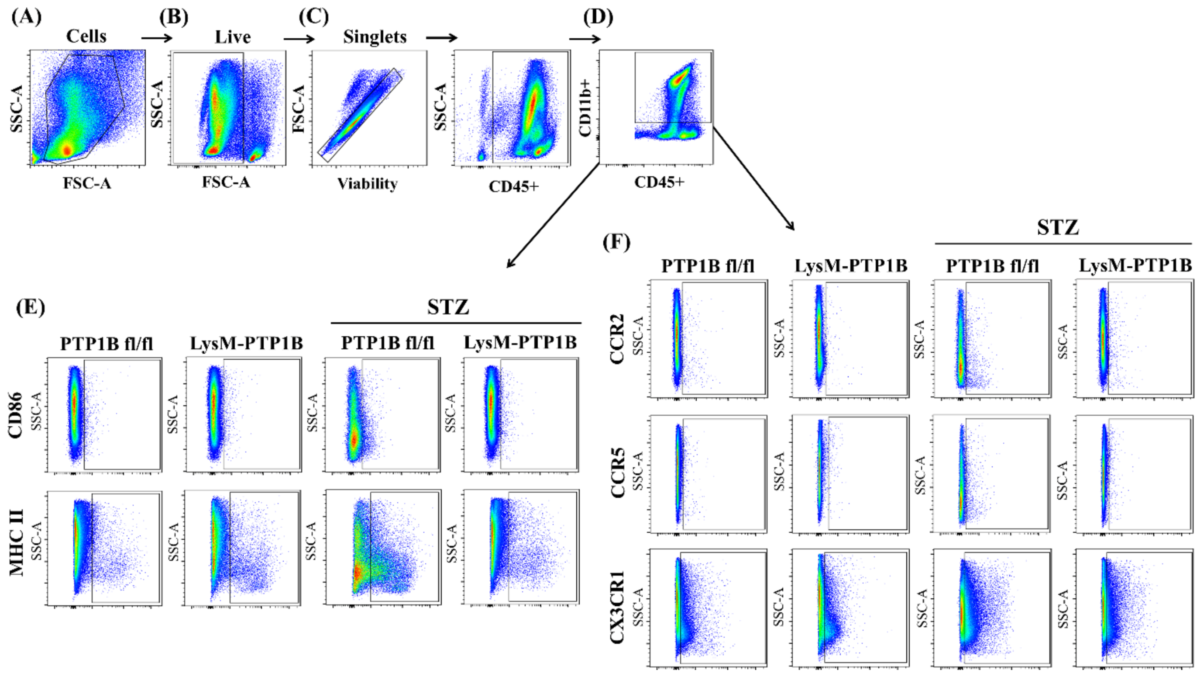


**Supplementary Figure 8. Gating strategy for flow cytometry of myeloid cell activation and chemokine receptors from BM and spleen tissues.** Single cell suspensions prepared from BM and spleen tissues of male and female mice at different time points were stained with pan-leukocyte marker (CD45), pan-myeloid marker (CD11b), immune activation markers (CD86 and MHC II) and chemokine receptors (CCR2, CCR5, and CX3CR1). Gating strategy was as follows; populations of leukocytes/cells were selected by forward vs side scatter gating (A). the live/dead cells were discriminated using eFluor 450 fixable viability dye (B). Single cells were selected within the live cells gate using forward scatter area vs forward scatter height (C). Populations of interest (CD45+ CD11b+) were selected within the CD45+-single cell-live cell-leukocyte gate (D). Representative dot plots of activation markers (CD86, MHC II) (E) and chemokine receptors (CCR2, CCR5, and CX3CR1) (F) expressing on the surface of CD45+CD11b+ cells in LysM-PTP1B and control littermate PTP1B-floxed mice with and without diabetes.

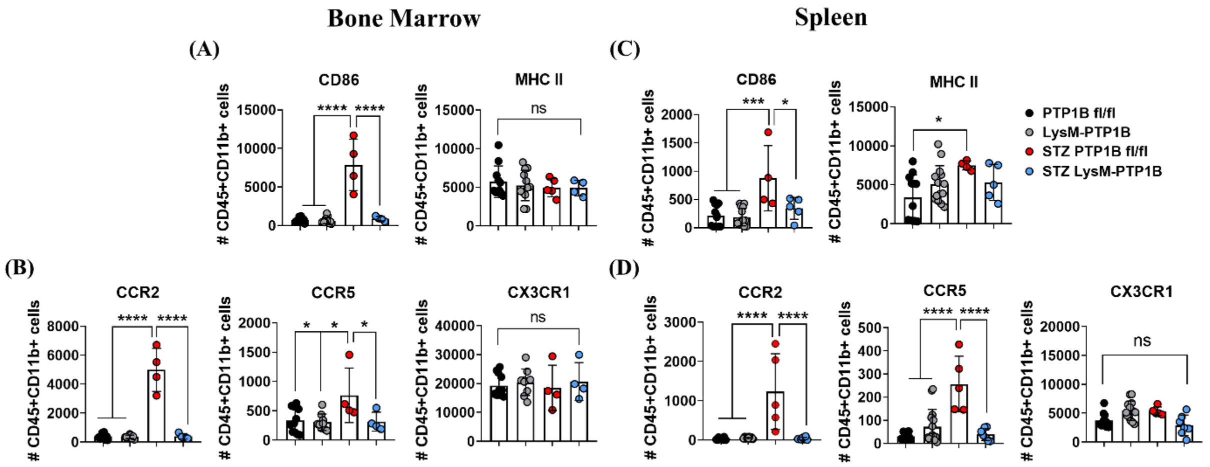


**Supplementary Figure 9. LysM-PTP1B express lower levels of myeloid cell activation in STZ mice.** Histogram scatter plots of absolute number of CD86 and MHC Class II expressing on CD45+CD11b+ cells from female mouse BM (A) and spleen (C). STZ-induced increased levels of CD86 and MHC Class II are decreased in LysM-PTP1B mice. Histogram scatter plots denote the absolute number of chemokine receptors from BM (B) and spleen (D) show similar decrease in chemokine receptor expression for CCR2 and CCR5 but not CX3CR1. Data are mean ± SD (n = 4 to 16 mice per group). One-way ANOVA was performed followed by Tukey's multiple comparison post-hoc test. (*P < 0.05; **P < 0.01; ***P < 0.001; ****P < 0.0001).

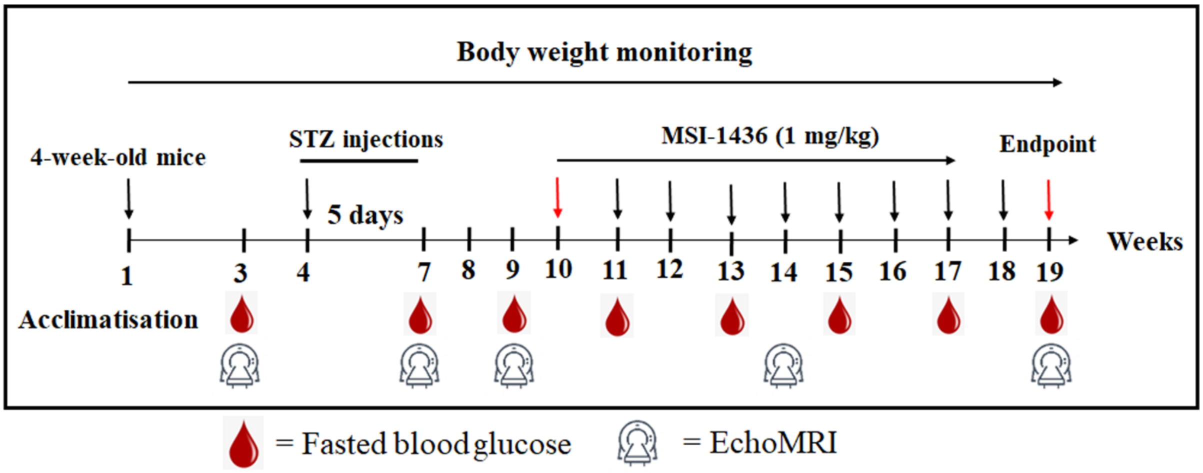


**Supplementary Figure 10. Experimental design of MSI-1436 treatment.** Schematic illustration of the experimental design for treatment of STZ-induced diabetic mice with MSI-1436 (1 mg/kg) once weekly for eight weeks.

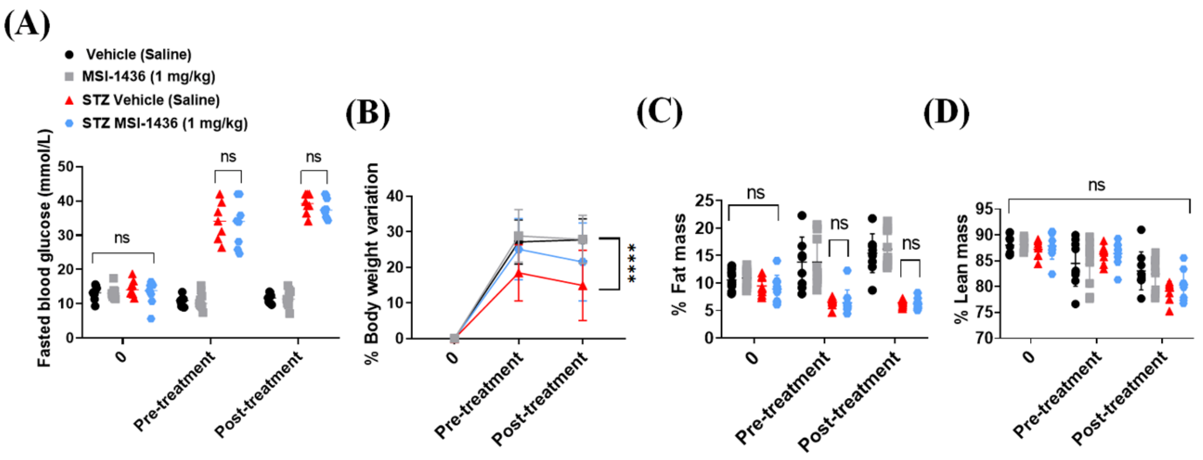


**Supplementary Figure 11. Fasted blood glucose, body weight and body composition are unchanged by MSI-1436 treatment in STZ mice.** Fasted blood glucose (A), body weight (B) and body composition (fat and lean mass) (C and D) were assessed in saline or MSI-1436 administered groups with and without diabetes. Values are shown as mean ± SD (n = 8 to 9 mice per group). Statistical significance determined by two-way ANOVA with Bonferroni's multiple comparison post-hoc test. (*$P < 0.05$; **$P < 0.01$; ***$P < 0.001$; ****$P < 0.0001$).

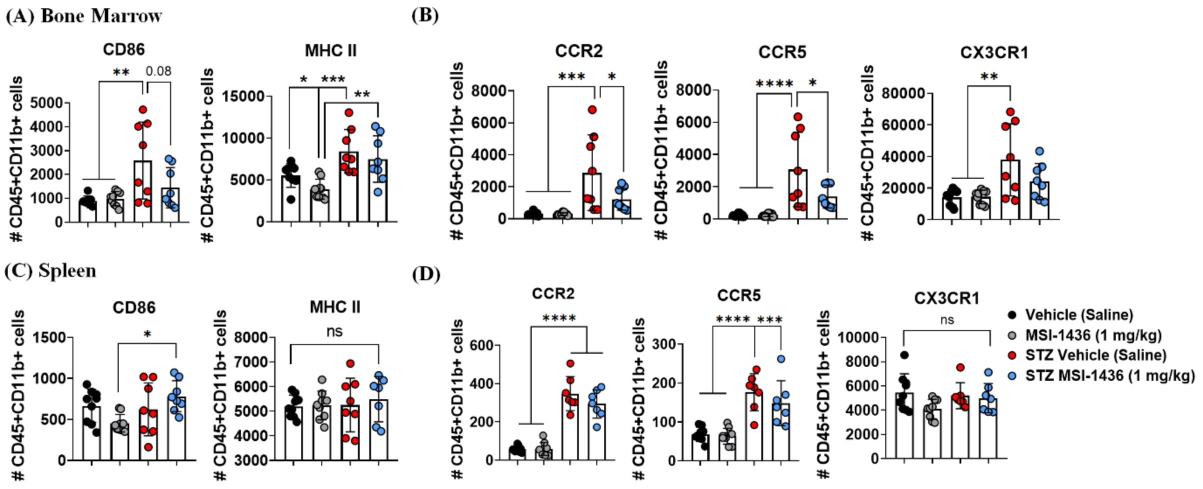


**Supplementary Figure 12. The small molecule PTP1B inhibitor, MSI-1436, prevented myeloid cell activation in STZ mice.** Scatter plots of absolute number of CD45+CD11b+ cells expressing activation and chemokine receptor markers from BM (A and B) and spleen (C and D) of saline or MSI-1436 treated mice with and without diabetes. Values are expressed as mean ± SD (n = 7 to 9 mice per group). One-way ANOVA was performed to indicate the statistical significance followed by Tukey's multiple comparison post-hoc test. (*$P < 0.05$; **$P < 0.01$; ***$P < 0.001$; ****$P < 0.0001$).
